# RHODOPSIN 7: An ancestral non-canonical photoreceptor shaping light-responsive behavior

**DOI:** 10.1101/2025.07.29.667052

**Authors:** Valentina Kirsch, Nils Reinhard, Heiko Hartlieb, Annika Mohr, Dirk Rieger, Peter Soba, Charlotte Helfrich-Förster, Pingkalai R. Senthilan

## Abstract

Animals rely on light not only for vision but also to adapt behavior to their environment through non-image-forming pathways. In *Drosophila*, most rhodopsins mediate retinal image formation, but RHODOPSIN 7 (RH7) is widely expressed in the brain and optic lobe, acting as a non-canonical light sensor reminiscent of mammalian melanopsin. We combined expression mapping, behavioral assays, and phylogenetic comparisons to investigate its function. RH7 detects temporal changes in luminance, guiding rapid behavioral responses, and regulating daily activity patterns. Flies lacking RH7 are less active during the dark phase, while strains with constitutively active RH7 show increased activity. Phylogenetic analyses suggest that RH7 is an ancient rhodopsin present across panarthropods, representing an intermediate between G protein-coupled receptors and specialized rhodopsins, highlighting a key step in the evolution of non-visual light perception. These findings show that RH7 functions as a melanopsin-like sensor, shaping behavior, and illuminating the origins of light detection.

## 2 Introduction

The ability to detect and respond to light is fundamental for many organisms and can influence behavior through both visual and non-visual pathways. Vision relies on specialized photoreceptor cells that convert light into neural signals through a process called phototransduction. This allows the brain to interpret visual information and guide behavior. Rhodopsin chromoproteins, which are composed of the seven-helix membrane apoprotein opsin and the chromophore retinal, play a central role in this process. When a photon is absorbed, the chromophore undergoes a conformational change, transforming from 11-cis-3-hydroxyretinal to all-trans-3-hydroxyretinal. Then, structural changes occur in the protein, initiating signal transduction. As G protein-coupled receptors (GPCRs), rhodopsins first activate a trimeric G protein, which then initiates a cascade of downstream signaling events in phototransduction (Xiong and Bellen, 2013).

While the primary function of rhodopsins is image formation in the visual system, some are also known to participate in non-image-forming functions. Based on their specific roles, rhodopsins are classified as visual or non-visual (Kingston and Cronin, 2016; Leung and Montell, 2017; Terakita, 2005). Several studies have highlighted the diverse functions of rhodopsins, including their involvement in thermosensation (Shen et al., 2011), hearing (Senthilan et al., 2012), proprioception (Zanini et al., 2018), and taste (Leung et al., 2020).

Opsins are classified into different groups, including ciliary opsins (c-opsins) and rhabdomeric opsins (r-opsins). While c-opsins trigger a G_alpha_ transducin (G_α_t)-mediated signaling pathway leading to Na^+^ channel closure and consequent hyperpolarization, r-opsins trigger a G_α_q-mediated depolarizing signaling cascade. In humans, c-opsins in rods and cones are essential for visual perception, while melanopsin (OPN4), an r-opsin expressed in intrinsically photosensitive retinal ganglion cells (ipRGCs), is involved in the synchronization of circadian rhythms and enhances contrast sensitivity (Sonoda et al., 2018). *Drosophila melanogaster* (*Dmel*) has seven rhodopsins (RH1-RH7), all belonging to the r-opsin group. While RH1-RH6 are expressed in well-characterized photoreceptors, the expression pattern and function of RH7 remain less clear and are still debated.

RH7 displays the canonical rhodopsin features, including seven transmembrane helices, a chromophore-binding lysine, and the LRTPXN motif located in the first intracellular loop (ICL1). However, two features mostly distinguish RH7 from other opsin groups: its extended N- and C-terminal tails, which are twice as long as in RH1-RH6, and the absence of the highly conserved QAKKMNV motif in the third intracellular loop (ICL3), important for G protein interaction (Figure 1A) (Senthilan and Helfrich-Förster, 2016). Additionally, RH7 contains a fully palindromic RCSI motif within its second intron, in contrast to the non-palindromic RCSI found upstream of other rhodopsins (Datta and Rister, 2022; Rister et al., 2015; Senthilan and Helfrich-Förster, 2016). The palindromic RCSI and its intronic location are likely key factors shaping the broader expression pattern of RH7. Tandem affinity purification of INTACT nuclei (TAPIN)-seq data indicate that *Rh7* is absent in retinal photoreceptors but present in visual processing neurons in the lamina and medulla (Figure 1B) (Davis et al., 2020).

**Figure 1.**
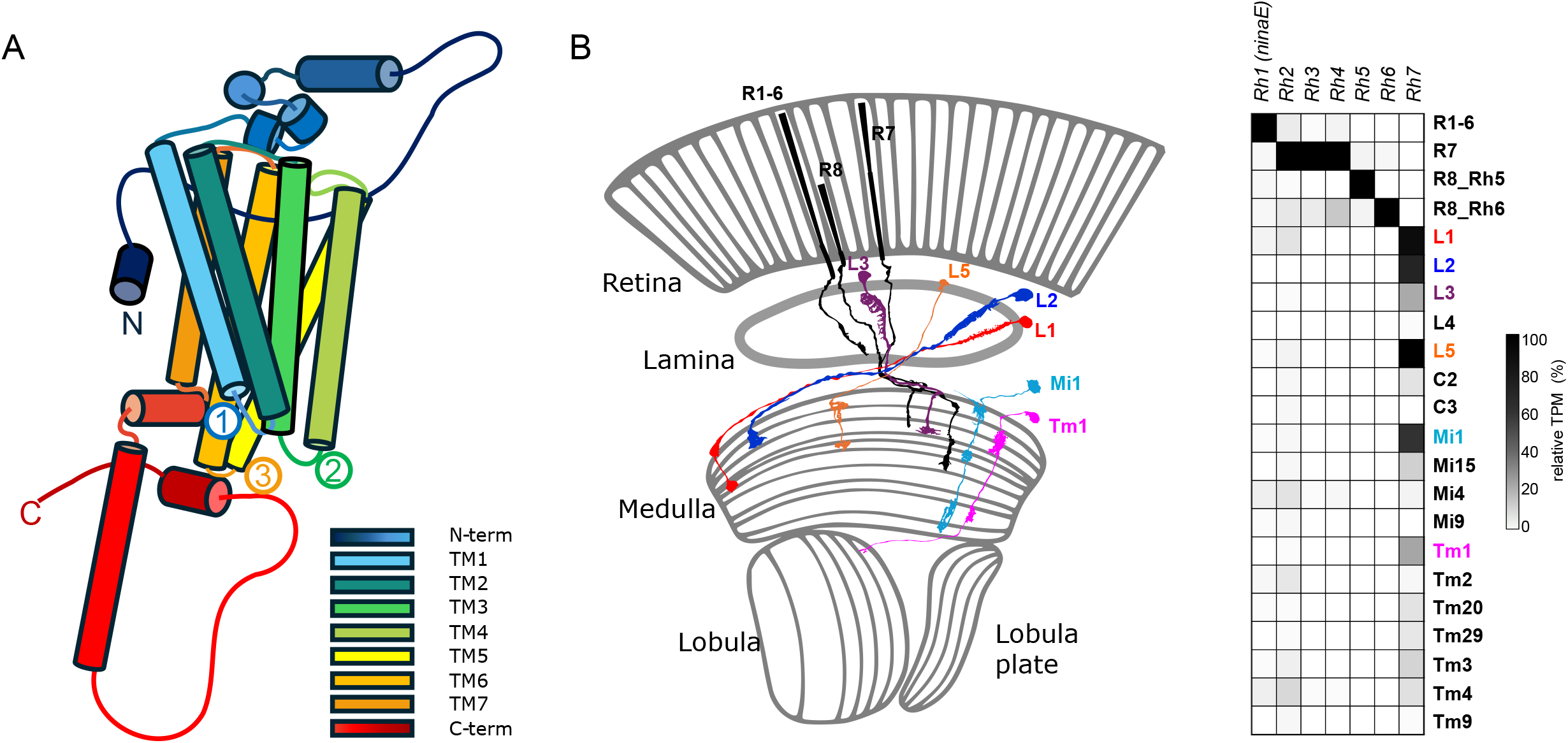
Predicted RH7 structure and expression pattern in the *Drosophila* optic system. **(A)** Schematic representation of the predicted 3D structure of RH7 (GH14208p) based on AlphaFold (Jumper et al., 2021). The structure is color-coded using a continuous spectral gradient ranging from blue (N-terminus) to red (C-terminus). The three intracellular loops are highlighted and labeled in numbered circles: (1) ICL1 with the LRTPXN motif, (2) ICL2 with the highly conserved DRY motif, and (3) ICL3, which lacks the typical QAKKMNV motif. **(B)** Left: Schematic representation of the *Drosophila melanogaster* optical system. The presumed *Rh7*-expressing cells, according to Davis et al, (2020) are shown (in color) in addition to retinal receptor cells (in black). Right: Heatmap of relative gene expression across *Drosophila* optic lobe cell types. TPM values were obtained by TAPIN sequencing (Davis et al., 2020) and normalized per gene (100% = maximum TPM per gene). Gray scale: from white (0%), to black (100%). Genes are shown on the x-axis, and cell types on the y-axis.

Previous studies on RH7 have yielded conflicting results regarding its function and expression, including discrepancies in the identified neuronal populations and associated behavioral phenotypes (Grebler et al., 2017; Kistenpfennig et al., 2017; Leung et al., 2020; Leung and Montell, 2017; Ni et al., 2017; Senthilan et al., 2019). Two factors may explain these discrepancies. First, RH7 is relatively large but likely adopts a compact, self-contained structure (Figure 1A). Its highly phosphorylatable N- and C-termini may interact with other proteins and further obscure relevant epitopes, making antibody detection nearly impossible. Second, the genetic background of the tested flies, particularly the presence of the *white* and *yellow* mutations as markers, appears to influence the *Rh7* phenotype, possibly due to overlapping expression sites. Both genes are expressed in the optic system (Davis et al., 2020) (Figure 3–figure supplement 2D) and are known to affect neuronal function and behavior (Bond et al., 2024; Drapeau et al., 2003; Rickle et al., 2025).

However, all previous studies, despite their differences, agree that *Rh7* is an atypical rhodopsin with non-visual functions, a hypothesis that is further supported by this study. We show that RH7 is distinct from other *Drosophila* rhodopsins. RH7 is predominantly expressed in optic lobe neurons and likely acts via a distinct signaling pathway, contributing to darkness detection and nocturnal activity. As an ancient rhodopsin, RH7 appears to contribute to basic visual functions, such as contrast processing as well as to non-visual functions like circadian synchronization, similar to melanopsins in mammals.

## 3 Material & Methods

### 3.1 Phylogenetics tree

Protein sequences were obtained from the NCBI RefSeq database (January 2026 annotation release, RS_2026_01). From these data, a dataset was compiled including all available invertebrate proteomes as well as selected vertebrate model organisms. Homologs of the opsins RH1, RH3, and RH7 were identified using BLASTP searches with an E-value cutoff of ≤ 1e−40. This procedure yielded sequences from 444 species. As *Paramacrobiotus metropolitanus* is currently the only non-arthropod panarthropod represented in RefSeq, *Ramazzottius varieornatus* and *Hypsibius exemplaris* were manually added to improve representation of Tardigrada, resulting in a final dataset of 446 species. All sequences in FASTA format are provided in Supplementary file 1.

To ensure balanced lineage representation in the phylogenetic tree, only one representative species per superfamily within each order was retained. Whenever possible, experimentally characterized species were preferentially selected as representatives. This resulted in 142 species with 558 protein sequences. Initial phylogenetic trees were examined to identify and remove obvious sequence duplicates, producing a final alignment of 470 protein sequences. Phylogenetic trees were constructed using Geneious Prime® (2026.0.2) with the Geneious Tree Builder. Protein sequences were globally aligned with free end gaps using the BLOSUM62 substitution matrix and gap costs of 10/0.1. Trees were used to assess evolutionary relationships among RH1, RH3, and RH7 opsins. Protein names, sequences, and accession numbers are provided in Supplementary file 1. To confirm phylogenetic assignments, reciprocal BLAST searches were performed against the *D. melanogaster* proteome, ensuring clade consistency and homology validation.

### 3.2 Co-occurrence analysis of RH7 and DCRY

*Drosophila*-type cryptochrome (DCRY) and RH7 homologs were identified from the RefSeq dataset. DCRY sequences were validated based on motif searches and cluster properties following the approach described in Deppisch et al. (2022), and RH7 homologs were validated based on the phylogenetic analysis performed in this study. All species containing homologs of RH1, RH3, or RH7 (E-value ≤ 1e−40) were included in the co-occurrence analysis. For species in which one of the proteins was detected but the other was not identified in RefSeq, additional searches were performed against clustered non-redundant (NR) protein datasets to detect potential DCRY or RH7 homologs missing from current RefSeq annotations.

### 3.3 GsX assay

To test the functionality of RH7, we used a GsX assay designed to determine which G_α_ proteins can bind to a given GPCR, in this case RH7 (Ballister et al., 2018). G protein coupling of GPCR constructs was examined using HEK293T or HEK293-ΔG7 cells, which lack the seven Gα subunits: G_α_s-type (GNAS, GNAL), Gαq-type (GNAQ, GNA11), Gα12/13-type (GNA12, GNA13), and G_α_z (GNAZ). All cells were obtained from A. Inoue at Tohoku University (Bowin et al., 2019). HEK293T cells were cultured in Dulbecco’s Modified Eagle Medium (DMEM) supplemented with 10% fetal bovine serum (FBS, Pan Biotech, Germany), 100 U/mL penicillin and 100 mg/mL streptomycin at 37 °C in 5% CO₂. The GsX assay was modified from Ballister et al. (Ballister et al., 2018). White 96-well plates (Greiner Bio-One) were coated with 0.1 mg/mL poly-L-lysine (Sigma Aldrich, USA) at 37 °C for 1 h prior to cell seeding. Receptor plasmids, G protein chimeras and the cAMP-sensitive luciferase reporter Glo22F (Promega) were co-transfected using Lipofectamine 2000 (Thermo Fisher, USA) and incubated at 37 °C in 5% CO_2_ for 24 hours. The medium was replaced with L-15 medium (phenol red-free, 1% FBS) containing 2 mM beetle luciferin (in 10 mM HEPES, pH 6.9) and 10 mM 9-cis-retinal or all-trans retinal and incubated for 1 hour at room temperature in the dark. cAMP-dependent luminescence was measured using a Berthold Mithras multimode plate reader (Berthold Tech., Germany). Cells were activated with a 1-second light pulse (470 nm) using an LED light plate (Phlox Corp., France) or CoolLED pE-4000 (CoolLED, UK). Two technical duplicates were performed, and data were normalized to pre-activation baselines.

### 3.4 Locomotor Activity

The locomotor activity of individual male flies was monitored at 1-minute intervals using the *Drosophila* Activity Monitoring (DAM) system (Trikinetics, Inc., Waltham, MA), which detects infrared (IR) beam interruptions. Male flies (3 to 7 days old) were individually placed in recording glass tubes (∼5 mm diameter) containing agar/sugar medium (2% agar; 4% sucrose) filled to one-third capacity and sealed with an air-permeable plug. Activity was quantified by counting the number of infrared (IR) beam crossings per minute in the center of each tube. All recordings were performed under constant conditions of 60% relative humidity and 25 °C. A standard day was defined as 12 h light (30 µW/cm^2^ ∼ 100 lux, the spectrum of the light is shown in Figure 3–figure supplement 1A) and 12 h darkness (LD 12:12). In addition, experiments using individual wavelengths (469 nm, 500 nm, and 521 nm) were performed; the corresponding absorption spectra are shown in Figure 3–figure supplement 1B. To assess light-off startle responses, which are brief clock-independent increases in activity after sudden light-off during the day, flies were repeatedly exposed to 30-minute dark pulses following 1.5 h illumination. Dark pulses were applied six times during each 12 h light phase. The experiments were conducted under “standard day” conditions consisting of 12 h light (30 µW/cm²) and 12 h darkness.

To account for differences in baseline activity between genotypes, the activity data were normalized to the mean activity level by dividing the activity values of each time point by the mean activity of the respective fly across the entire 24 h recording period. For all experimental conditions, data from four consecutive days were averaged, omitting the first day to reduce effects of acclimatization.

In a second startle response assay (4-hour assay), we specifically aimed to assess the acute startle response independent of circadian influences. To minimize contributions of the circadian clock, flies were first rendered arrhythmic by exposure to constant light (120 µW/cm², ∼500 lux) for two days. This led to sustained activation of the blue light receptor DCRY and continuous degradation of TIMELESS, resulting in a standstill of the molecular clock (Deppisch et al., 2022; Dolezelova et al., 2007; Emery et al., 2000). Subsequently, flies were subjected to three consecutive days of 4-hour light-dark cycles (6 cycles per day). Each cycle consisted of 3.5 hours of light (5 µW/cm² (∼10 lux), 30 µW/cm² (∼100 lux) or 120 µW/cm² (∼500 lux)) followed by a 30-minute dark phase (no light or reduced light intensity of 5 µW/ cm² or 30 µW/ cm²). This short-cycle paradigm prevents re-entrainment of circadian rhythms and allows isolation of the startle response itself (see schematic representation in Figure 3–figure supplement 1C). Startle activity was recorded during each cycle. The first 4 h cycle was excluded from analysis to avoid carryover effects from the prior light regime. Values from all remaining 17 cycles were averaged for analysis. Data were plotted with the 30-minute dark pulse centered on the time axis, resulting in two light periods of 105 minutes each, before and after the dark pulse. For normalization, the mean activity during the 105 minutes preceding light-off was set to one, and activity after light-off was expressed relative to this baseline.

### 3.5 Locomotor activity analysis

Activity data were analyzed in R using custom scripts. Raw activity traces were smoothed using cubic smoothing splines (smooth.spline, default settings). Morning and evening activity peaks were identified as the maximal smoothed activity within predefined windows (morning: 330–390 min; evening: 1050–1110 min). Peak amplitude was calculated as the mean activity across the peak maximum ±2 min.

Activity onset and offset were detected from the smoothed signal using a threshold defined as 80% of the mean daily activity. An onset was defined when activity levels were first for at least 10 min below the threshold followed by at least 10 min of activity above the threshold together with a positive signal derivative; offsets were defined analogously using inverse criteria. Search windows were morning onset (180–360 min), morning offset (420–720 min), evening onset (750–1080 min), and evening offset (1080–1200 min). Daytime activity (avg activity day) was quantified as the summed smoothed activity from the light phase (361–1080 min). Siesta beginning time was defined as the morning offset. Siesta length was calculated as the time interval between morning offset and evening onset. Activity after lights-off (LOFF activity (min)) was calculated as the interval between the evening offset and lights-off (evening offset − 1080). Maximum activity after lights-off (LOFF max activity) was defined as the highest activity value recorded after lights-off, expressed as the maximum number of infrared beam crossings per minute.

### 3.6 Fly lines, Primers, and Antibodies

The lines listed below were used for the experiments in this study. To minimize background effects, all lines were previously brought on the same X chromosomal background derived from the control fly CantonS (CS), carrying wild-type alleles of the *yellow* (*y⁺*) and *white* (*w⁺*) genes, meaning any mutations in these genes are absent.

### 3.7 Generating the *Rh7-Gal4* line

We generated a *Rh7-Gal4* line to investigate *Rh7* expression. To this end, the *Rh7* promoter was cloned upstream of a *Gal4* coding sequence, driving *Gal4* expression in *Rh7*-expressing cells. Bioinformatic analyses suggested that the *Rh7* promoter is unusual and relatively large (Senthilan and Helfrich-Förster, 2016), hence we used a region including 1,080 bp upstream of exon 1 and extending into exon 2, which contains the translation start site. Due to the overall length of the target region (>8,000 bp), the construct was assembled in four fragments, which were sequentially cloned into the *pPTGAL* vector (Sharma et al., 2002). The four inserts were amplified from *D. melanogaster* genomic DNA using specific *Gal4* primer pairs (see Table 2). Insert 1 and insert 4 included artificial restriction sites (Eco52 and KpnI, respectively), while all other restriction sites were naturally occurring. Cloning began with insert 4 (see Figure 4–figure supplement 1) and proceeded stepwise with the remaining fragments. Final plasmids were amplified in *E. coli* NEB 10-beta and the construct integrity was confirmed by sequencing. The plasmid was injected into *w^1118^* flies by TheBestGene Inc., and the resulting transgenic line was crossed into the same *y^+^w^+^* background as the controls (Table 1).

**Table 1.**
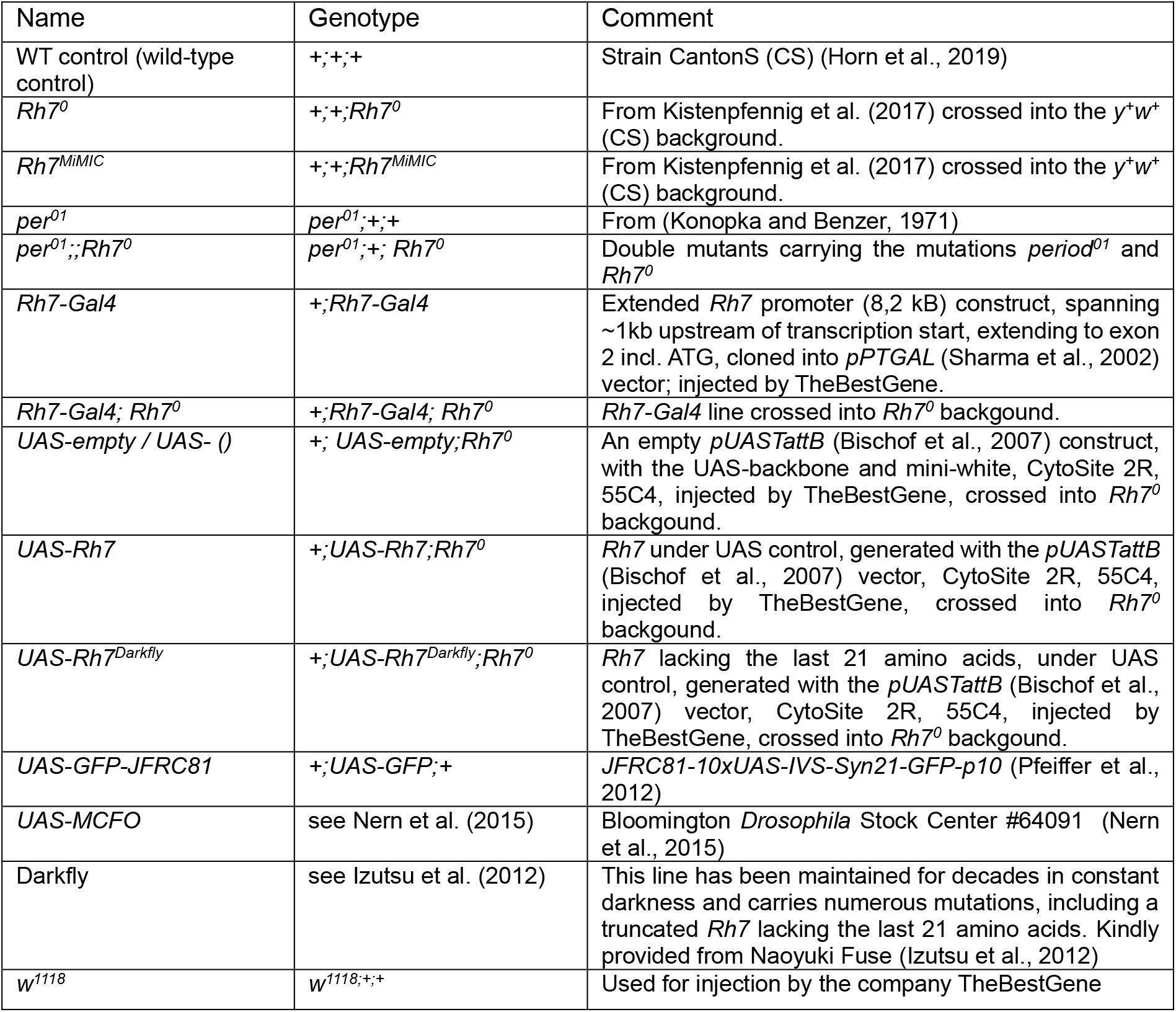
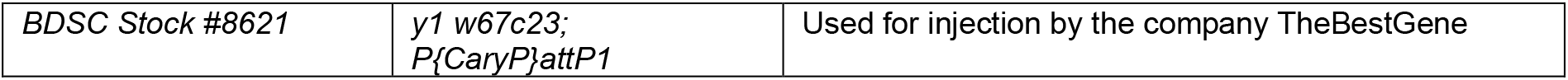
Fly lines.

**Table 2.**
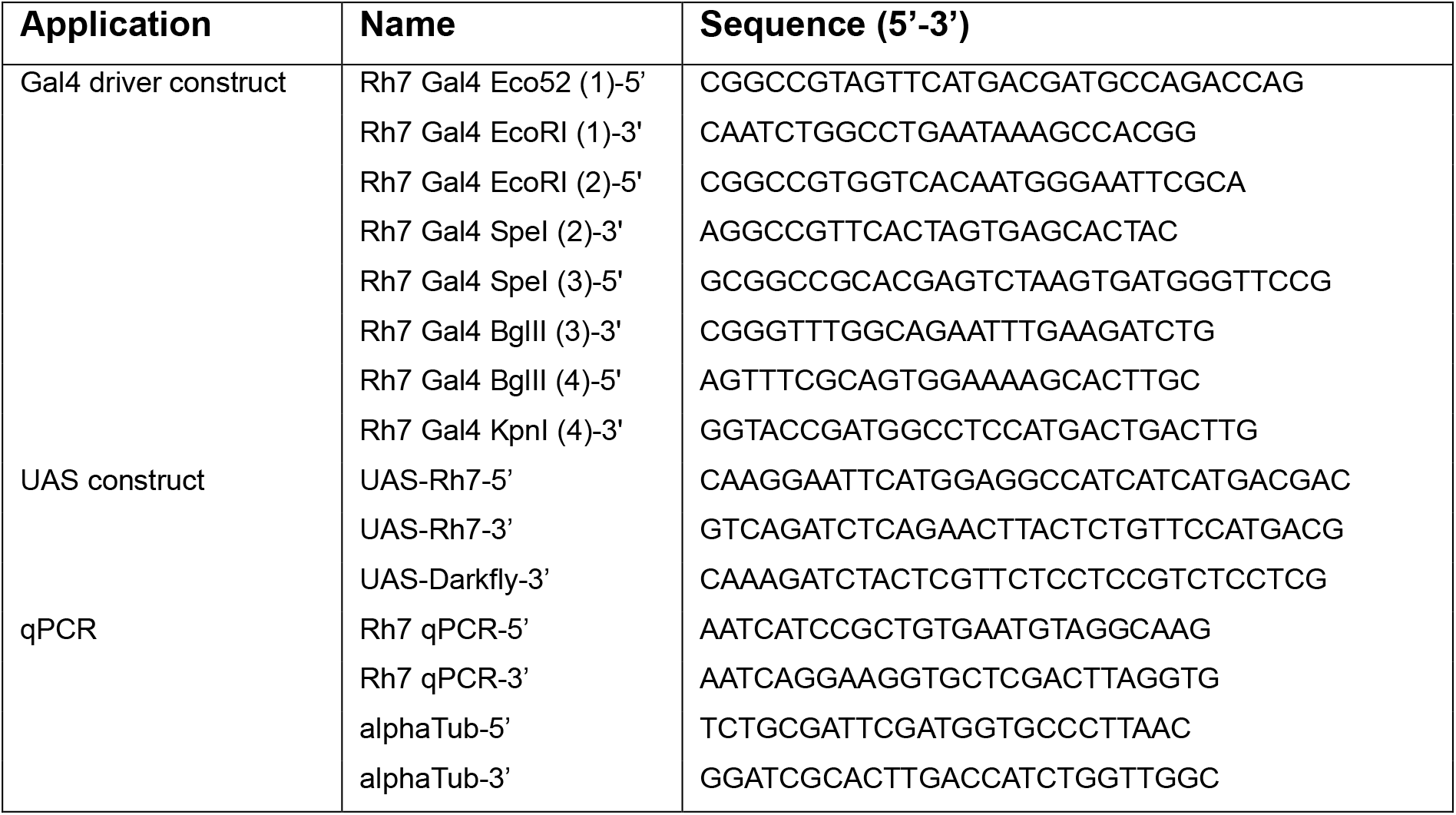
Primers.

### 3.8 Generating UAS-constructs

For *Rh7*-rescue experiments, UAS constructs were generated to assess the ability of different *Rh7* variants to rescue the phenotype. *UAS-Rh7* carries the wild-type *Rh7* sequence, *UAS-Rh7^Darkfly^* carries the Darkfly variant, and *UAS-empty* (*UAS-())* serves as a negative control (no-rescue). The *Rh7* construct was prepared from the *D. melanogaster* cDNA using the primers UAS-Rh7-5’ and UAS-Rh7-3’ (Table 2), digested with EcoRI and BglII enzymes and ligated into the *pUASTattB* vector (Bischof et al., 2007). For the *UAS-empty* construct, the *pUASTattB* vector was used unmodified. The vectors were then injected into flies with CytoSite 2R, 55C4 (BDSC Stock #8621, containing attP1) by the company TheBestGene Inc. All lines were brought into a *+;+;Rh7^0^* background and finally crossed with flies carrying respective *Gal4*-constructs in *+;+;Rh7^0^* background and assayed. The *UAS-Rh7^Darkfly^* construct was prepared with the primers UAS-Rh7-5’ and UAS-Darkfly-3’ (Table 2) with the same steps described above. All transgenic lines were crossed into the same *y^+^w^+^* background as the controls (Table 1).

### 3.9 Optomotor response

To investigate the contribution of RH7 in the lamina and medulla to visual motion detection, male flies were individually tested for their optomotor walking behavior. Flies were raised under standard LD 12:12 conditions at 25 °C. Subsequently, they were maintained under constant light (LL, 120 µW/cm^2^ (∼500 lux)) for at least 3 days to induce arrhythmicity and minimize the influence of the circadian clock on the optomotor response. The optomotor response setup comprises: (1) an upright cylinder (ø 8 cm; H 4.5 cm) with vertical black and white stripes (2.2 cm width and 4 cm height) inside for visual stimulation and (2) a transparent Plexiglas arena (ø 3 cm; H 1.5 cm) with a tube for inserting the flies. The open space available for movement was only about 2 mm, limiting the flies to a single horizontal walking plane (Figure 5–figure supplement 1C). The LED-illuminated cylinder with 6 white and 6 black stripes rotated clockwise (CW) and counterclockwise (CCW) at 16 rpm, equivalent to a frequency of 1.6 Hz. The flies were acclimatized for 2 minutes without light, followed by 2 minutes under CW rotation with light, a 1-minute rest without light, and then 2 minutes under CCW rotation with light. For the optomotor response (OR), the number of complete laps around the arena and complete body turns in the direction of the pattern movement within both 2-minute runs were added, while movements in the opposite direction were subtracted. The result was then divided by the total number of cylinder rotations (4×16). A representative video of the optomotor response assay is provided as Supplementary Video 1.

### 3.10 qPCR

For quantitative PCR (qPCR), total RNA was extracted from either five heads, five brains, 10 retinae or 10 laminae of control flies. Flies were raised under standard LD 12:12 conditions at 25 °C. Tissues were carefully dissected from adult flies without prior fixation to avoid interference with RNA integrity and downstream extraction procedures. Dissections were performed during the light phase and were not restricted to specific time points. Total RNA was extracted using the Quick-RNA™ Microprep Kit (Zymo), and cDNA was synthesized with the QuantiTect Reverse Transcription Kit (Qiagen). For qPCR, three biological replicates were used for each tissue, with two technical replicates. qPCRs were performed using the SensiFAST™ SYBR® No-ROX Kit (Bioline) in the Qiagen Rotor-Gene Q qPCR machine, with raw data fit by dynamic tube and slope correction calculation using the Rotor-Gene Q Series software. Relative mRNA levels were calculated using the Δ-CT equation, with *α-tubulin* used as a reference gene (ΔCT = CT_α-*tubulin*_ − CT*_Rh7_*). qPCR was performed with *Rh7* (Rh7 qPCR-5’, Rh7 qPCR-3’) and control (alphaTub-5’, alphaTub-3’) primers, each at 0.1 mM (Table 2).

### 3.11 Immunohistochemistry

Fluorescent immunohistochemistry was performed on 3 to 5 days old male flies. Whole flies were fixed for 3 h in 4% paraformaldehyde in 0.5% PBST (1xPBS with 0.5%Triton X-100) followed by three 10-minute rinses with PBS. The brains were dissected in 0.5% PBST and incubated overnight at 4 °C with 5% normal goat serum (NGS) or normal donkey serum (NDS) and 0.02% NaN_3_ in 0.5% PBST, depending on the host species of the secondary antibodies. Antibody solutions contained the antibody in the concentration specified in Table 3 and 5% NGS or NDS with 0.02% NaN_3_ in 0.5% PBST. The brains were incubated for two days at 4 °C in primary antibody solution followed by five 10-minute washes with 0.5% PBST. Subsequently, the brains were incubated for at least 4 hours at room temperature or overnight at 4 °C with the secondary antibody solution containing the corresponding secondary antibodies (Table 3). The samples were washed five times for 10 minutes with 0.5% PBST before embedding in VectaShield 1000 mounting medium (Vector Laboratories, Burlingame, CA, USA) using object slides with spacers (∼170 µm). The high-precision coverslips (170 ± 5 µm, No.1.5H) were sealed with FixoGum. Brains with retinae attached were embedded with retinae facing the object slide to ensure easy imaging from the posterior. Object slides were stored in the dark at 4 °C until scanning. In addition, individual neurons were visualized using the MultiColor FlpOut (MCFO) technique (Nern et al., 2015), which stochastically labels single cells with distinct membrane-targeted fluorescent tags. Flippase-mediated excision of FRT-flanked stop cassettes enables sparse and color-resolved labeling, allowing detailed analysis of neuronal shape and projections. Labeled brains were processed and mounted as described above.

**Table 3.**
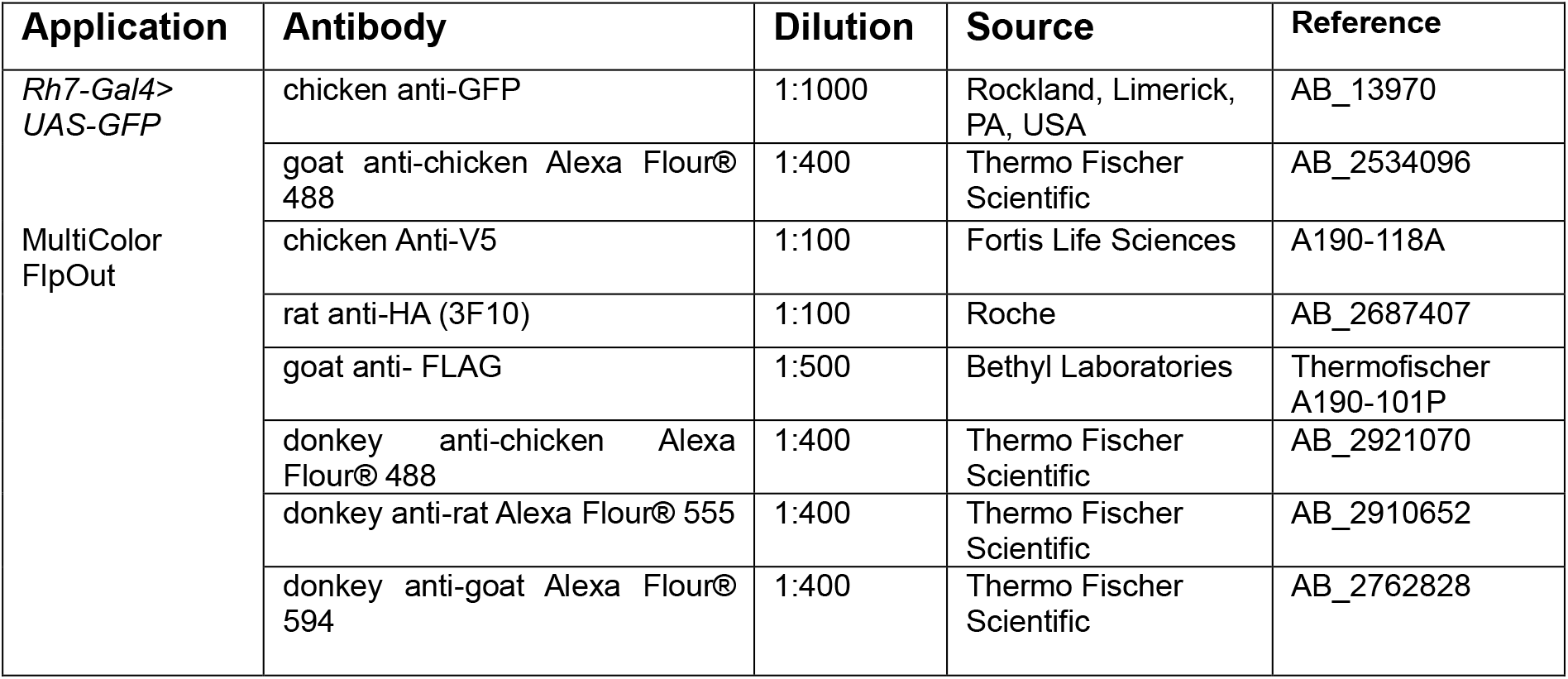
Antibodies used for immunohistochemistry.

### 3.12 *In situ* hybridization

For RNA *in situ* hybridization, we used the hybridization chain reaction v3 (HCR) method developed by Molecular Instruments (Choi et al., 2018). The flies were fixed and dissected as described for the fluorescent immunohistochemistry. The tissue was either used directly for *in situ* hybridization or dehydrated and stored in 99% methanol at −20 °C for up to 14 days. Dehydration was performed by sequential incubation in 30%, 50%, 70%, and 90% methanol in PBS (2 minutes each). Rehydration was performed in the opposite order. *In situ* HCR was performed using a modified protocol from Bruce et al. (2021). In short, *Rh7* probe mixes containing 20 probe pairs were designed by Molecular Instruments. Instead of using Proteinase K or detergent-based denaturation solutions, tissues were incubated in 0.5% PBST for 30 minutes. This was followed by a 30-minute incubation at 37 °C in preheated probe hybridization buffer (Molecular Instruments), and then an overnight incubation (≥ 20 h) at 37 °C in preheated probe hybridization buffer containing the *Rh7* probe mix (12 nM). After hybridization, the tissue was washed four times for 15 minutes each with preheated probe washing buffer (Molecular instruments) at 37 °C. This was followed by two washes at room temperature (5 minutes each) in 5xSSCT (5xSSC buffer with 1% Tween20). Next, the tissues were incubated for 30 minutes at room temperature in amplification buffer (Molecular Instruments) followed by incubation for at least 20 h at room temperature in amplification buffer containing the hairpins required for signal amplification. Hairpins were prepared by heating 3 µM h1 hairpins and 3 µM h2 hairpins separately in amplification buffer to 95 °C for 90 seconds. They were then cooled to room temperature for 30 minutes, protected from light, before being mixed and applied to the tissue. When hairpins were reused, the already mixed hairpins were reheated together. After five washes with 5xSSCT (2 x 5 minutes, 3 x 30 minutes) the tissues were either directly embedded in VectaShield as described above, or the normal fluorescent immunohistochemistry protocol was used for additional antibody labeling starting with blocking in 5% NGS or NDS.

### 3.13 Image acquisition and analysis

Images were acquired with a Leica TCS SP8 confocal microscope equipped with a photon multiplier tube and hybrid detector. A white light laser (Leica Microsystems, Wetzlar, Germany) was used for excitation. We used a 20-fold glycerol immersion objective (0.73 NA, HC PL APO, Leica Microsystems, Wetzlar Germany) for wholemount scans. We obtained 8-bit confocal stacks with 2048 × 1024 pixels with a maximal voxel size of 0.3 x 0.3 x 2 μm and an optical section thickness of 3.12 μm. Weak signals were amplified using line accumulation. Images were analyzed using Fiji. No changes in pixel intensity were applied other than linear changes in brightness and contrast.

## 4 Results

### 4.1 Evolutionary position and distribution of RH7

To determine the taxonomic distribution of RH7 among r-opsins, a phylogenetic tree was constructed. The analysis clearly identifies the three main clades of *Drosophila* rhodopsins: clade 1 (RH1, RH2, and RH6), clade 2 (RH3, RH4, and RH5), and clade 3, which corresponds to RH7. Unlike the first two clades, which include only rhodopsin homologs from other arthropods, the RH7 clade also contains homologs from tardigrades, forming a distinct branch within the clade (Figure 2A).

**Figure 2.**
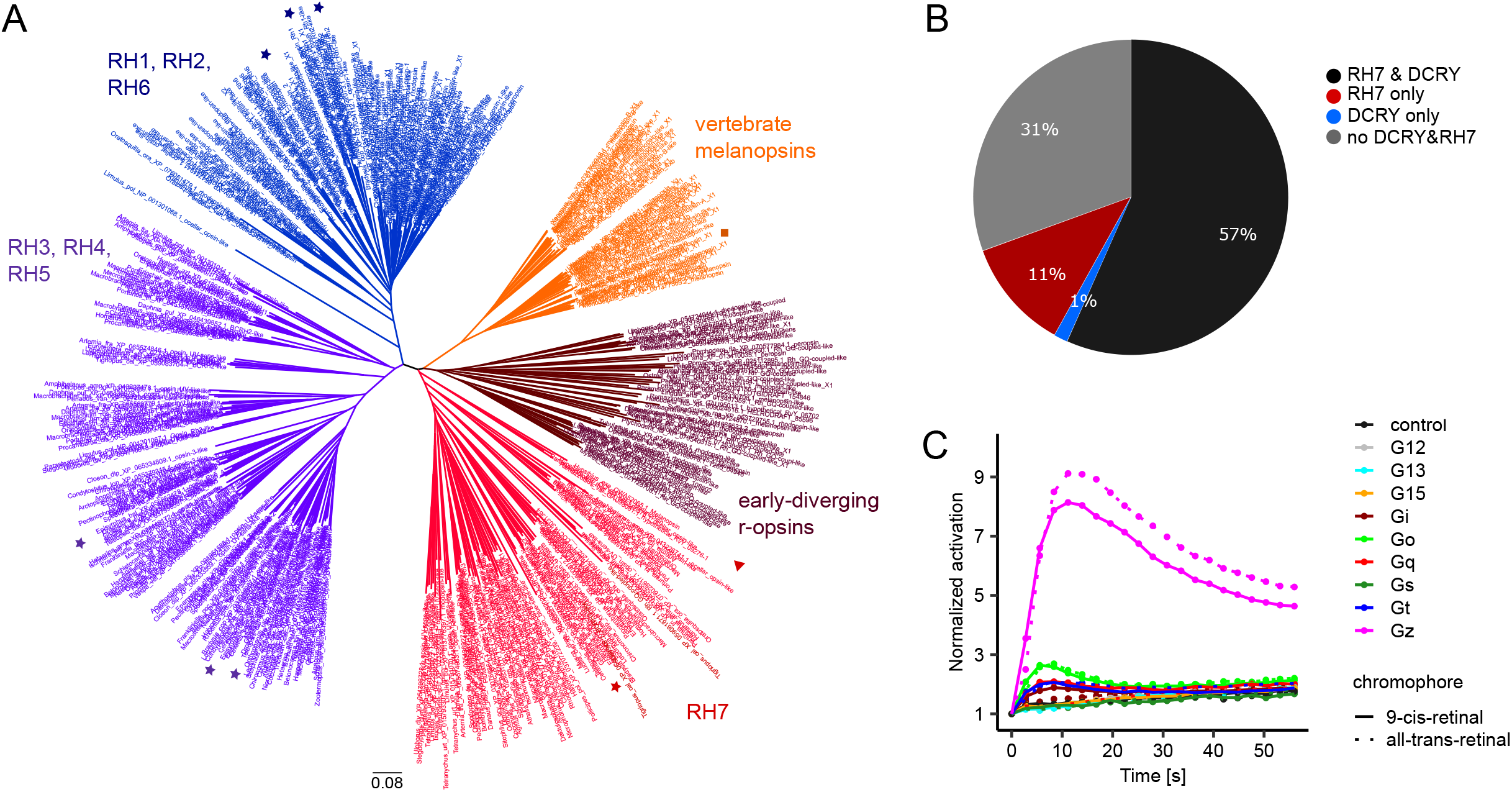
Evolutionary conservation, distribution, and signaling properties of RH7. **(A)** Phylogenetic tree of 470 r-opsin protein sequences from 142 selected species, highlighting the clades containing the *Drosophila* rhodopsins: RH1, RH2, and RH6 (dark blue); RH3, RH4, and RH5 (purple); and RH7 (red). Early-diverging r-opsins are shown in brown, and vertebrate melanopsins in orange. *Drosophila* opsins are marked with stars in the corresponding colors. The four tardigrade RH7-like opsins are represented with a red triangle, and the human melanopsin is indicated by an orange square. **(B)** Distribution of RH7 and DCRY across 360 analyzed panarthropod species: 57% possess both genes (black), 12% possess only RH7 (red), 1% possess only DCRY (blue), and 32% lack both proteins (grey). **(C)** GsX assay showing that RH7 robustly activates the G_α_z chimera in a light-dependent manner, with minimal differences between 9-cis-retinal and all-trans-retinal. No activation was detected with the G_α_q chimera.

Adjacent branches contain more ancestral r-opsins from a broad range of metazoans, including mollusks, annelids, arthropods, and echinoderms. One early-diverging branch, highlighted in orange, corresponds to the vertebrate melanopsin clade, which itself appears to have split into two subgroups early in its evolution. While mammals have lost one of these lineages, all other vertebrates retain both. The placement of RH7 near the mammalian melanopsin clade and other early-diverging r-opsins suggests that it occupies an ancestral position within the r-opsin family. To assess the distribution of DCRY and RH7, we examined all species available in the RefSeq dataset, totaling 446 species. DCRY is widely distributed across Bilateria, whereas RH7 arose later within Panarthropoda. Among 360 panarthropod species, 58% possess a DCRY homolog and 68% an RH7 homolog. Overall, 57% of species contain both genes, 1% only DCRY, 11% only RH7, and 31% neither (Figure 2B). Nearly all species with DCRY (∼98%) also retain RH7. Conversely, 17% of RH7-containing species lack DCRY. This pattern is observed across several insect orders, including basal hymenopterans of Tenthredinoidea, coleopterans, hemipterans, and dipterans. It is also seen in other arthropod lineages such as chelicerates, tardigrades, and some crustaceans. Together, these observations highlight the strong co-occurrence of RH7 and DCRY, as well as the existence of lineages where RH7 persists without DCRY, suggesting potential functional and evolutionary relationships between these photoreceptors.

### 4.2 Functional distinction of RH7 from canonical *Drosophila* opsins

We investigated RH7 signaling by examining its interaction with different G_α_ proteins using a GsX assay (Ballister et al., 2018). This assay employs chimeric G_α_ proteins that share a core derived from G_α_s, which triggers cAMP production upon activation. The C terminal 14 amino acids of each chimera, critical for GPCR binding, are replaced with sequences from other G_α_ subtypes, such as G_α_q or G_α_z. This allows each chimera to mimic the coupling specificity of a distinct G protein while ensuring that any productive GPCR interaction results in a measurable increase in cAMP levels. For this analysis, we co-expressed RH7 with various GsX chimeras in HEK cells and stimulated RH7 with brief light pulses. We observed robust activation with the G_α_z chimera, with both 9-cis-retinal and all-trans-retinal (Figure 2C). Although there is no G_α_z homolog in *Drosophila*, related G_α_ subunits such as G_α_o and G_α_i are expressed in the fly. Nevertheless, these results revealed two key findings: (1) RH7 can be activated by light, and (2) RH7 does not appear to signal via the G_α_q-mediated pathway, unlike other *Drosophila* rhodopsins. These findings indicate that RH7 exhibits a signaling profile distinct from other insect rhodopsins.

### 4.3 *Rh7^0^*mutants show altered daily activity pattern & reduced light-off startle responses

To study the influence of RH7 on the daily locomotor activity of flies, wild-type (WT) control flies and *Rh7^0^* mutants were examined under standard LD 12:12 conditions (Figure 3A). During the day, mutants and wild-type controls showed the morning and evening activity typical of *Drosophila melanogaster* with a siesta in between. However, *Rh7^0^* mutants showed a significant increase in daytime activity compared to WT controls. In addition, mutants displayed a significantly shortened siesta, which also started later than in controls. Another prominent difference was observed in the startle response after light-off, characterized by a transient increase in locomotor activity lasting approximately 5–10 minutes. While WT control flies showed a pronounced startle response after lights-off, this response was strongly reduced in *Rh7^0^* mutants and terminated more rapidly after lights-off. To determine whether these behavioral differences depended on the spectral composition of the light, locomotor activity was additionally analyzed using distinct wavelengths (Figure 3–figure supplement 1B). Overall, locomotor activity patterns were comparable across the different wavelength conditions, and the differences between WT and *Rh7^0^* mutants remained evident.

**Figure 3.**
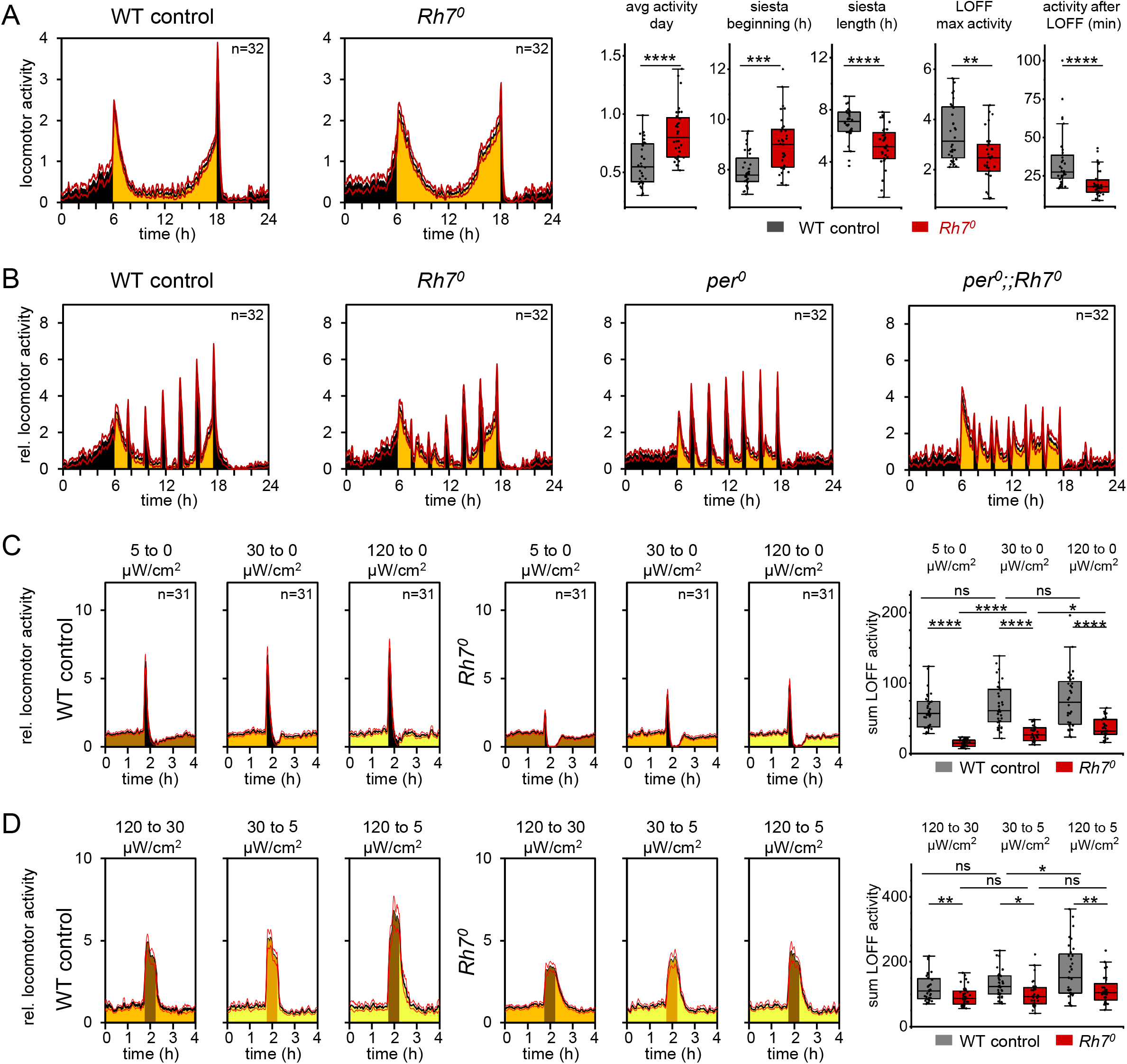
Altered locomotor activity and light-off responses in *Rh7^0^* mutant flies. **(A)** Left, average locomotor activity profiles of wild-type (WT) and *Rh7^0^* mutant flies under standard conditions (12 h light (30 µW/cm²):12 h dark, 25 °C). Right, box plots showing daytime average activity, siesta onset, siesta duration, maximum activity after lights-off (LOFF), and LOFF response duration. **(B)** Average activity profiles of WT, *Rh7^0^, per^0^,* and *per^0^;;Rh7^0^* flies during a 12 h light (30 µW/cm²):12 h dark cycle at 25 °C. During the light phase, the light was switched off for 30 min every 1.5 h to assess the light-off startle response. Activity was normalized to the mean activity of each fly across the entire 24 h recording period. Non-normalized activity profiles are shown in Figure 3–figure supplement 2A. **(C)** Light-off startle responses after 3.5 h of light exposure at 5, 30, or 120 µW/cm², followed by a 30 min dark period. **(D)** 4h assay with dimmed light. After 3.5 h of light exposure, illumination was reduced for 30 min to either 5 or 30 µW/cm² instead of being switched off completely (****p <0.0001, ***p <0.001, **p <0.01, *p <0.05, Mann-Whitney U-Test, BH correction, R-studio).

### 4.4 Startle response to lights-off persists without a functional clock

The startle response, triggered by the sudden light-off, differs from circadian-regulated evening activity and can be triggered during the day as well. To test the startle response, we performed 30-minute dark pulses every 1.5 hours during the light phase on a standard LD 12:12 day and recorded the fly’s activity. We observed that the startle responses of both WT controls and *Rh7^0^* mutants were generally lower in the morning than in the afternoon (Figure 3B). The circadian clock and thus the internal state of flies appear to modulate the startle response, as can be seen in clockless *per^01^* mutants. *per^01^* mutants exhibited a startle response that remains constant throughout the day. This startle response is significantly and consistently reduced in the *per^01^;+;Rh7^0^* double mutants (Figure 3B). These results show that the startle response is modulated by the circadian clock but is not dependent on it, and that RH7 plays an important role in this response.

In a second assay (4-hour assay), flies were rendered arrhythmic by continuous exposure to bright light to isolate the startle response from circadian influences. We then tested how different light intensities affected this response using repeated 4-h cycles over 3 days, consisting of a 3.5 h light phase at different intensities (5, 30, or 120 µW/cm²) and a 30-minute dark phase. Smaller intensity changes were assessed by replacing the dark phase with dim light (5 or 30 µW/cm²), while the light phase remained brighter (30 or 120 µW/cm², respectively). Under all tested conditions, *Rh7^0^* mutants exhibited a significantly lower startle response than WT controls. Both genotypes increased the startle response with increasing light intensity, but this reached only significance for *Rh7^0^* mutants (Figure 3C). *Rh7^MiMIC^* mutants, an independent *Rh7* mutant line retaining reduced but detectable *Rh7* mRNA levels, displayed a similar phenotype to *Rh7*^0^ mutants (Figure 3–figure supplement 2B, 2C). The phenotype persisted in trans-heterozygous *Rh7^0^/Rh7^MiMIC^* flies but was absent in heterozygous controls, further supporting a role of RH7 in mediating the startle response (Figure 3–figure supplement 2E, 2F).

To investigate whether the startle response is also induced by reductions in light intensity rather than transitions to complete darkness, we performed the same assay while maintaining residual light levels of 5 or 30 µW/cm² during the “dark phase”. Transitions to complete darkness induced a pronounced, short-lasting startle response, whereas transitions to residual-light conditions resulted in sustained elevated activity throughout the 30-minute stimulus without a sharp transient peak (Figure 3D). Across all conditions, WT flies showed the strongest responses. *Rh7^0^* mutants displayed similar qualitative response patterns, but with a markedly reduced magnitude compared to WT controls (Figure 3D). In WT flies, larger differences in light intensity tended to evoke stronger responses. In contrast, *Rh7^0^* mutants showed largely unchanged response levels across the different light conditions.

### 4.5 *Rh7* is expressed in the optic lobe and clock neurons

To determine the tissue-specific expression of *Rh7*, we performed qPCR analyses using *α-tubulin* as a reference gene. The expression of *Rh7* was highest in the lamina, followed by the brain, while only minimal levels were detected in the retina (Figure 4A). Additionally, we performed *Rh7* RNA *in situ* hybridization chain reaction (HCR), which verified the strongest *Rh7* expression in the lamina and medulla (Figure 4B). We further observed considerably weaker expressions in neurons of the central brain (Figure 4B). No *in situ* hybridization was detected in *Rh7^0^* mutants (Figure 4C).

**Figure 4.**
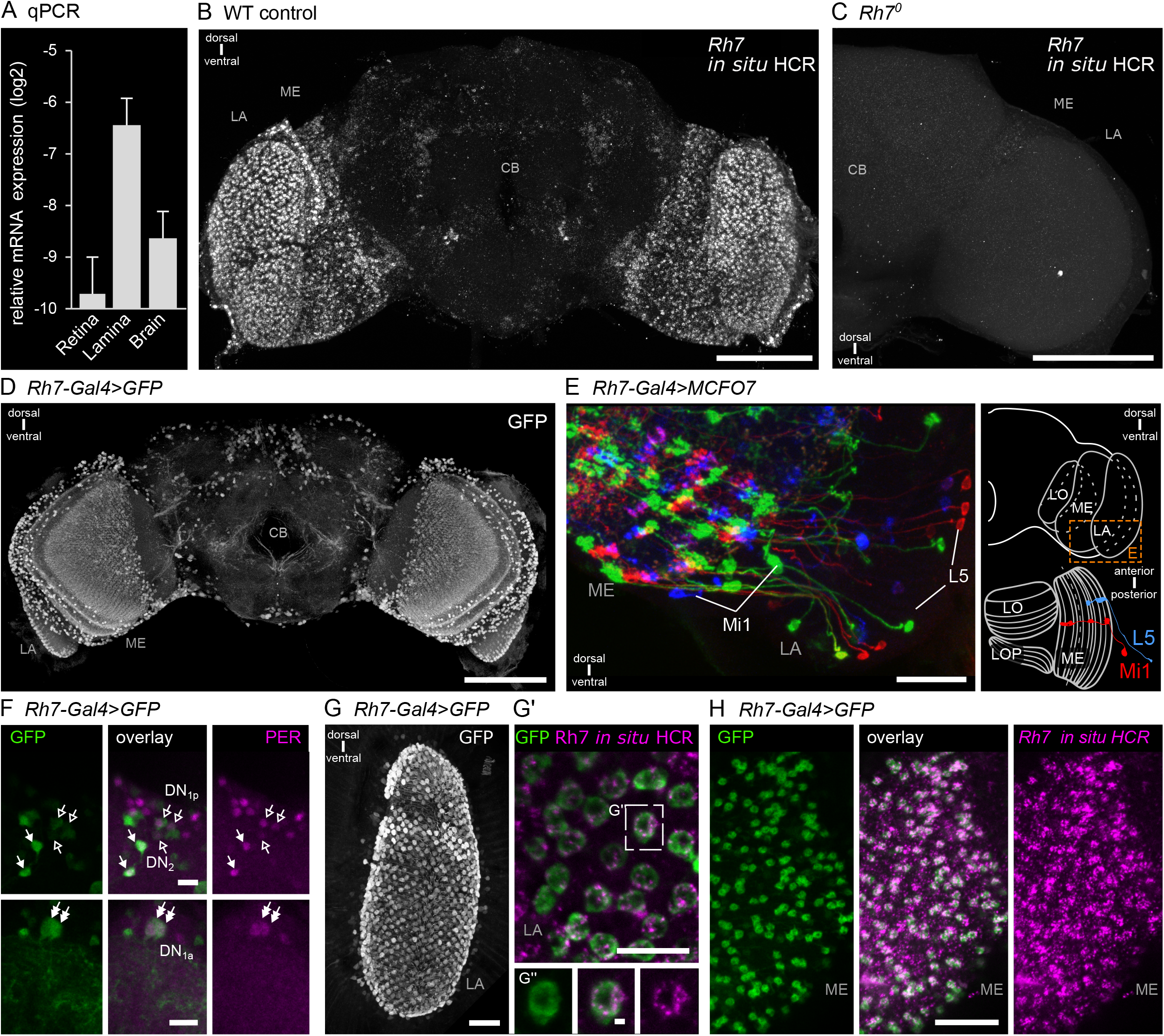
Expression pattern and cellular localization of *Rh7* in the optic lobe and circadian neurons. **(A)** Relative qPCR analysis revealed strong *Rh7* expression in the lamina, followed by the brain, with minimal background expression in the retina. **(B)** *Rh7 in situ* HCR in wild-type (WT) controls revealed strong *Rh7* signals in the lamina (LA) and medulla (ME), and weaker signals in the central brain (CB). **(C)** *Rh7^0^* mutants showed no detectable *Rh7 in situ* HCR signal, confirming the specificity of the probe and the loss of *Rh7* expression in the mutant. **(D)** *Rh7-Gal4*-driven *UAS-GFP* expression revealed robust labeling in the optic system, particularly in the lamina and medulla, as well as in various brain cells. **(E)** *Rh7-Gal4*-driven MCFO labeling identified lamina neurons L5 and the medulla neuron Mi1. Right: schematic illustration of the detected neurons in the optic lobe. **(F)** Immunostaining of *Rh7-Gal4>UAS-GFP* flies with anti-GFP and anti-PER antibodies revealed co-labeling in clock neurons, suggesting that a subset of circadian neurons expresses *Rh7*. Open arrows mark DN_1p_ and arrows mark DN2; double arrows mark DN_1a_. **(G)** & **(H)** *in situ* HCR in *Rh7-Gal4>UAS-GFP* flies revealed that all GFP-labeled cells in the lamina **(G)** and medulla **(H)** also showed *Rh7* transcript signal, confirming *Rh7* expression in these neurons. Scale bar in B, C, D represents 100 µm, in E, G, H 25 µm, in F, G’ 10 µm and in G’’ 1 µm.

To further investigate *Rh7* expression, we generated a *Rh7* promoter-driven *Gal4* line (*Rh7*-*Gal4*). Since the *Rh7* gene expression regulatory elements are not only located within the promoter but also extend into the transcriptional region, we cloned a large part of this region, including exon 1, intron 1 and part of exon 2, together with the start codon (Figure 4– figure supplement 1). When comparing GFP expression driven by the newly generated *Rh7-Gal4* with the *Rh7* RNA *in situ* hybridization and qPCR data, we again observed the strongest expression in the lamina and medulla. Interestingly, also the *Rh7-Gal4* line showed additional expression in neurons of the central brain (Figure 4D). To further characterize *Rh7*-expressing neurons, we used MultiColor FlpOut (MCFO) (Nern et al., 2015), which stochastically labels neurons when crossed to a *Gal4*-driver line. Using MCFO, we identified at least two cell types of the optic system based on their characteristic arborization pattern in the medulla: lamina neuron 5 (L5) and medulla intrinsic neuron 1 Mi1 (Figure 4E). Importantly, both neurons were also present in the TAPIN RNA-seq analysis, with L5 showing the strongest *Rh7* expression (Davis et al., 2020). Next, we used an antibody against the clock protein PERIOD to test for colocalization, given that previous studies showed *Rh7* expression in subsets of clock neurons (Kistenpfennig et al., 2017; Ma et al., 2021; Ni et al., 2017). We observed colocalization of GFP and PER in some DN_1p_, DN_1a_, and DN_2_ neurons (Figure 4F) but not in the LN_v_s contrary to Ni et al. (2017). To test whether *Rh7* expressing clock neurons also express *dcry,* we performed simultaneous RNA *in situ* hybridization for *dcry and Rh7.* As suggested by single cell RNA sequencing data, the clock neurons expressing *Rh7* are CRY-negative with the DN_1a_ forming the only exception (Figure 4– figure supplement 1B).

To validate the expression pattern of the *Rh7-Gal4* line, we performed *in situ* hybridization with *Rh7-Gal4>GFP* flies. All GFP-labeled cells were also detected by RNA *in situ* hybridization, although the signal was less pronounced in cells of the central brain. Some cells, particularly in the optic system, were detected only by *in situ* hybridization and not by the *Gal4* line. This suggests an even broader expression of *Rh7* than shown by the *Gal4* line (Figure 4G, 4H).

### 4.6 C-terminal truncation of RH7 affects dark-phase behavior

The so-called Darkfly was kept in complete darkness for 57 years (1400 generations) accumulating mutations that may have enhanced its adaptation to perpetual darkness (Izutsu et al., 2012). Notably, the mutant carries a nonsense mutation in the *Rh7* gene, resulting in the loss of the final 21 amino acids at the C-terminus. qPCR and immunohistochemistry analyses indicate preserved *Rh7* transcript levels and expression patterns, suggesting that the truncated RH7 protein is indeed expressed (Figure 3–figure supplement 2C, Figure 5–figure supplement 1B). When we exposed the Darkfly to dark pulses in a 12:12 light-dark cycle, they exhibited the opposite phenotype to *Rh7^0^* mutants, showing a stronger startle response (Figure 5A), possibly reflecting RHODOPSIN 7 adopting a constitutively active form in the Darkfly strain, driving the enhanced behavioral response (Figure 5B).

**Figure 5.**
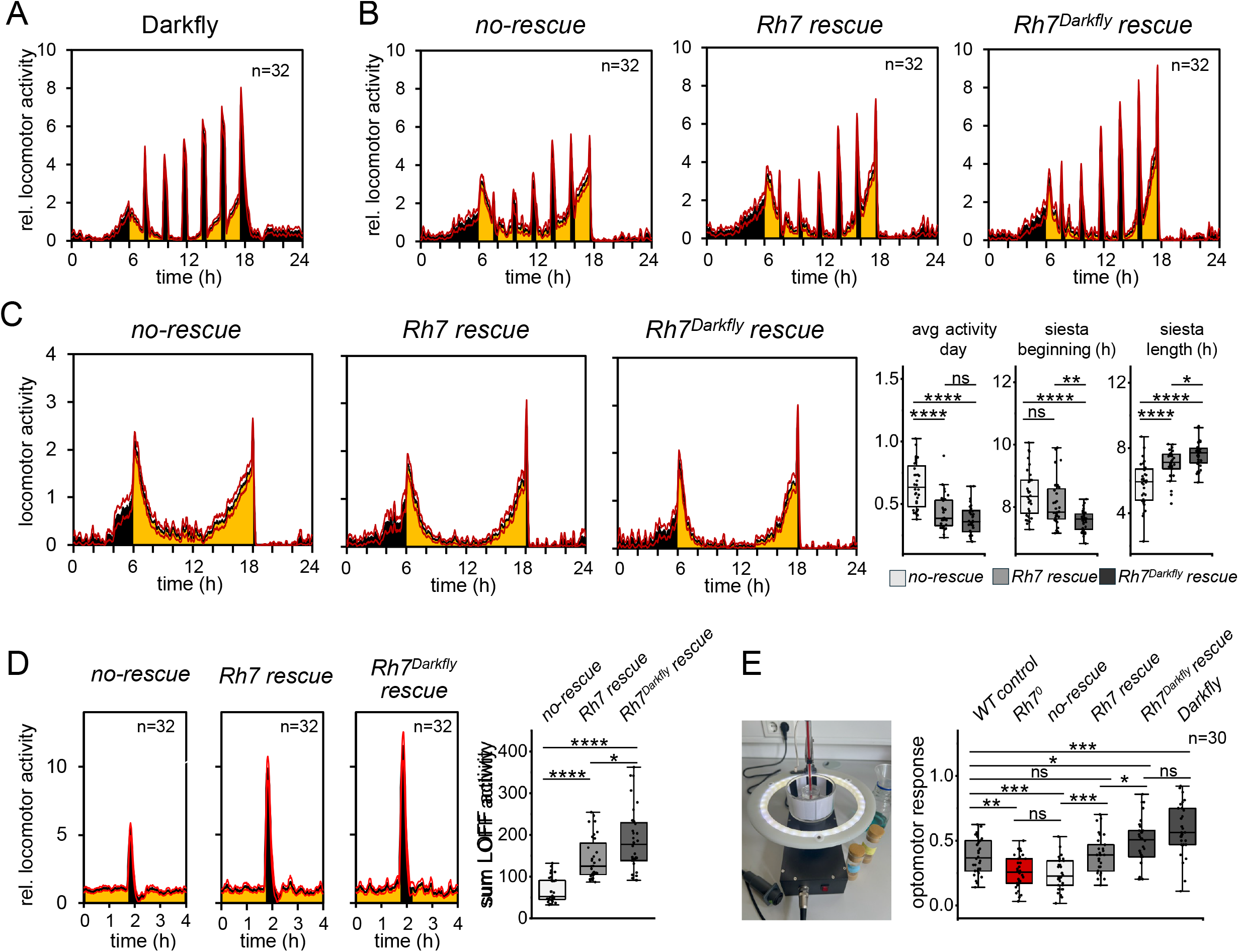
Functional characterization of *Rh7* rescue and Darkfly rescue lines. **(A)** & **(B)** Normalized average daily activity of Darkfly and rescue fly lines recorded under standard conditions (12 h light (30 µW/cm²):12 h dark, 25 °C). During the light phase, lights were turned off for 30 min every 1.5 h. **(A)** Darkfly flies exhibit an enhanced startle response to light-off. **(B)** No-rescue (*Rh7-Gal4; Rh7^0^>UAS-empty; Rh7⁰*) display a phenotype similar to *Rh7* mutants, whereas the *Rh7* rescue line (*Rh7-Gal4; Rh7^0^>UAS-Rh7; Rh7⁰*) resembles WT controls. The *Rh7^Darkfly^* (*Rh7-Gal4; Rh7^0^*>*UAS-Rh7^Darkfly^; Rh7⁰*) rescue shows an enhanced startle response similar to Darkfly, particularly during the second half of the day. **(C)** RH7 rescue alters daytime activity and siesta parameters under LD conditions. Both *Rh7* rescue and *Rh7^Darkfly^* rescue flies showed reduced average daytime activity compared to no-rescue flies. Similar effects were observed for siesta length, including a significant difference between *Rh7* rescue and *Rh7^Darkfly^* rescue flies. For siesta beginning (h), significant differences were observed between no-rescue fliesand *Rh7^Darkfly^* rescue flies, as well as between *Rh7* rescue and *Rh7^Darkfly^* rescue flies**. (D)** To measure the startle response, the 4-hour assay was performed. The *Rh7* rescue line restored the enhanced startle response that was absent in the no-rescue mutants, which behave like *Rh7⁰* mutants (p < 0.0001). The *Rh7^Darkfly^* rescue showed an even stronger startle response, significantly higher than the *Rh7* rescue line (p < 0.001). **(E)** Optomotor response assays show that no-rescue flies behave like *Rh7⁰* mutants, with no significant difference between them. *Rh7* rescue flies respond like WT controls. Both *Rh7⁰* mutants and no-rescue flies differ significantly from WT and *Rh7* rescue flies (p < 0.01). Darkfly and *Rh7^Darkfly^* rescue flies do not differ from each other but show significant differences compared to WT and *Rh7* rescue flies (***p < 0.001, **p < 0.01, *p < 0.05, Mann-Whitney U-Test, BH correction, R-studio).

To validate the role of RH7 in the startle response, we conducted a rescue experiment by expressing *Rh7* in *Rh7^0^* mutants using the *UAS-Rh7* construct driven by *Rh7-Gal4*. As a control for cloning and insertion artifacts, we used a *UAS-empty* line with the same insertion site as *UAS-Rh7*, driven by *Rh7-Gal4* in the *Rh7^0^* background. In the dark pulse experiment, *Rh7-Gal4>UAS-empty;Rh7^0^* (“no-rescue”) flies behaved like mutants, while *Rh7-Gal4>UAS-Rh7;Rh7^0^* flies (“*Rh7* rescue”) resembled WT controls (Figure 5B). To express a truncated RH7 protein lacking the last 21 amino acids, as found in the Darkfly, we crossed the *UAS-Rh7^Darkfly^; Rh7^0^* line with the *Rh7-Gal4; Rh7^0^* line (“*Rh7^Darkfly^* rescue”). We observed an increased startle response in the offspring, most prominently during the second half of the light phase (Figure 5B). Under standard LD 12:12 conditions, both *Rh7* rescue and *Rh7^Darkfly^* rescue flies showed reduced average daytime activity compared to no-rescue flies. Similar effects were observed for siesta length, with *Rh7^Darkfly^* rescue flies additionally differing significantly from *Rh7* rescue flies. Differences in siesta beginning were also detected, with significant changes observed between no-rescue flies and *Rh7^Darkfly^*rescue flies, as well as between *Rh7* rescue and *Rh7^Darkfly^* rescue flies (Figure 5C). In the 4-hour assay, in which the flies were made arrhythmic we found a clear rescue of the startle response in *Rh7-Gal4>UAS-Rh7;Rh7^0^* flies compared to the *Rh7-Gal4>UAS-empty;Rh7^0^*controls. Furthermore, *Rh7-Gal4>UAS-Rh7^Darkfly^;Rh7^0^*flies showed a significantly increased startle response (Figure 5C).

### 4.7 RH7 modulates optomotor responses

The *Drosophila* optomotor response is a reflex that stabilizes locomotion by counteracting perceived environmental motion with its own movement. Motion detection relies on the processing of spatiotemporal changes in luminance across the visual field. The optic system, particularly the lamina neurons, plays an important role in this response (Yang and Clandinin, 2018). We found that *Rh7^0^* mutants show deficits in the optomotor response, while the Darkfly showed an increased optomotor response (Figure 5D). When the rescue flies were tested, the *Rh7-Gal4>UAS-empty;Rh7^0^*flies behaved like the *Rh7^0^* mutants, whereas the *Rh7-Gal4>UAS-Rh7;Rh7^0^* flies showed a significant rescue, with no significant difference from WT controls. In contrast, *Rh7-Gal4>UAS-Rh7^Darkfly^;Rh7^0^* flies showed a significantly higher optomotor response than the WT controls and the *Rh7-Gal4>UAS-Rh7;Rh7^0^*flies. This response was similar to the Darkfly (Figure 5E).

## 5 Discussion

### 5.1 RH7 is an ancestral and structurally distinct opsin

Although RH7 has been known for nearly 25 years, its function remains poorly understood. This is partly due to its atypical expression outside the retina and its divergence from canonical visual rhodopsins. Phylogenetic analyses suggest that RH7 is an early-diverging r-opsin within Panarthropoda. Homologs identified in arthropods and tardigrades (Figure 2A) indicate a deep evolutionary origin and place RH7 within an ancient opsin lineage. Neighboring branches include early-diverging r-opsins from lophotrochozoans and echinoderms, as well as melanopsins from vertebrates. This phylogenetic position supports the ancestral character of RH7 and suggests that it may retain features of early opsin function, as reflected by RH7-related opsins in tardigrades, which have been linked to monochromatic vision (Fleming et al., 2021; Hering and Mayer, 2014). Many species that possess RH7 inhabit sun-exposed environments, suggesting that RH7 may contribute to light sensitivity under natural conditions. As our results with different light contrast and the optomotor tests suggest, Rh7 may enhance contrast sensitivity, which is advantageous for insects when they move rapidly through environments with highly variable light contrast, for example, under trees between shaded and well-lit areas.

The ancestral character of RH7 is further reflected in its unusual structural features with long N- and C-terminal regions carrying a high density of predicted phosphorylation sites. This suggests a high potential for post-translational regulation and possibly increased signaling flexibility. The extended termini distinguish RH7 from the more compact structure of typical rhodopsins and may represent an intermediate stage between canonical GPCRs and specialized visual rhodopsins (Figure 6A). In addition, RH7 lacks several conserved motifs found in other *Drosophila* rhodopsins, which likely emerged during the specialization of visual opsins. The absence of these features, together with the extended termini, supports the idea that RH7 occupies a structurally and evolutionarily distinct position. A more detailed comparison of these motifs and their functional implications is presented in section 5.5.

**Figure 6.**
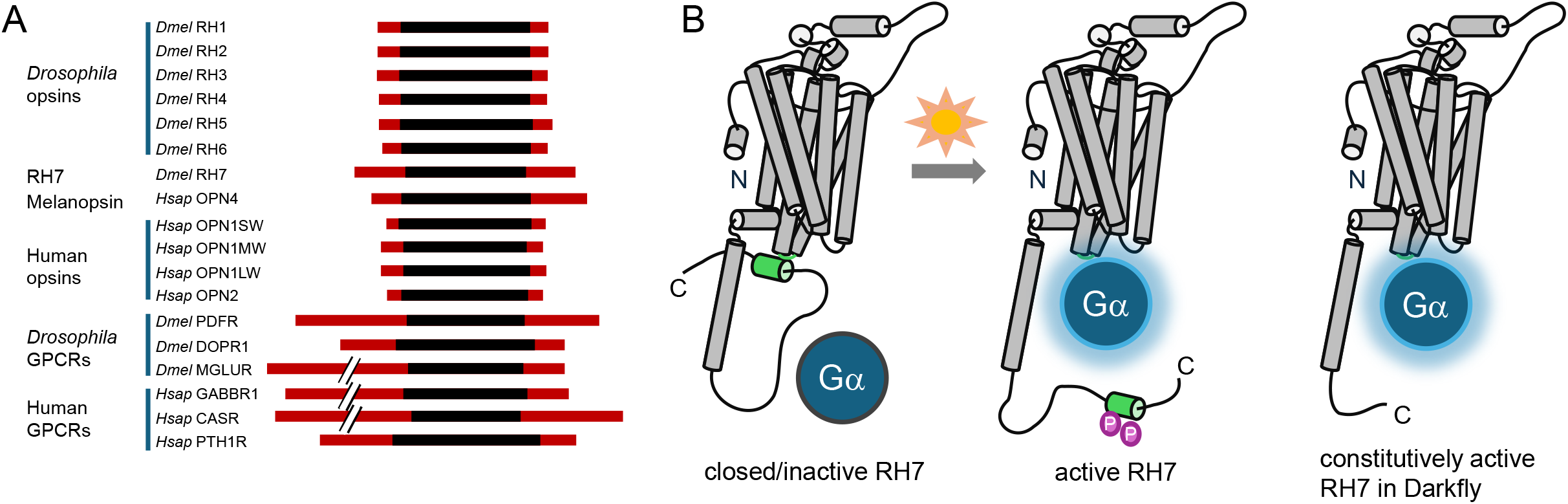
Structural features and hypothetical activation states of RH7. **(A)** Protein lengths of *Drosophila* and human opsins are shown alongside randomly selected non-opsin GPCRs. General GPCRs are significantly longer, primarily due to extended N- and C-termini, whereas opsins are shorter and largely confined to the transmembrane core. RH7 and OPN4 (melanopsin) display intermediate protein lengths relative to both groups. The red bars represent the N- and C-termini of the proteins, and the black bars represent the core protein, spanning from the 1st to the 7th transmembrane domain. Additional information, including full protein names and UniProt IDs, is provided in Supplementary table 1. **(B)** Left: Predicted 3D structure of RH7 based on AlphaFold, representing a closed conformation. Center: A putative open conformation that may result from light-induced conformational changes and subsequent C-terminal phosphorylation (magenta), potentially promoting G protein binding, as suggested by Maggio et al. (2023). Right: Hypothetical structure of RH7 in the Darkfly variant, possibly stabilized in an open, constitutively active state, independent of light.

Besides its structural divergence, the developmental origin and central localization of RH7-expressing neurons support its ancestral photoreceptive role. RH7 is expressed in central visual neuropils such as the lamina and medulla that process input from peripheral photoreceptors. Unlike canonical r-opsins, which are primarily expressed in specialized retinal photoreceptor cells, RH7 expression in central brain regions suggests an ancient form of non-image-forming photoreception integrated directly within neural circuits. This idea is further supported by the presence of opsins expressed in brain regions of other early-diverging bilaterians, such as annelids and mollusks (Arendt et al., 2004; Vöcking et al., 2015; Wollesen et al., 2019).

While our data show *Rh7* expression in the optic lobe and central brain, previous work also reports atypical expression in retinal support cells and multidendritic neurons (Charlton-Perkins et al., 2017; Lazopulo et al., 2019). We did not detect signals in these cells, likely due to low expression levels and background fluorescence. However, the low *Rh7* mRNA levels detected by qPCR (Figure 4A), together with the faint staining reported by Kistenpfennig et al. (2017), may indicate *Rh7* expression in retinal support cells. Overall, these findings are consistent with our observations and further highlight the non-canonical expression of RH7. RH7 thus likely represents a central, evolutionarily conserved opsin for light sensing, predating the split between sensory input and processing centers.

### 5.2 Divergence from canonical signaling: first insights into RH7 phototransduction

The signaling properties of RH7 were explored using heterologous GsX assays, which clearly indicated that RH7 can be activated by light (Figure 2C) but does not interact with G_α_q. This lack of interaction appears plausible, as G_α_q expression is restricted to retinal cells, whereas *Rh7* is predominantly expressed in lamina and medulla neurons (Davis et al., 2020). Instead, RH7 appears to interact with mammalian G_α_z. Although *Drosophila* lacks a direct homolog of G_α_z, the related inhibitory subunits G_α_i and G_α_o from the same family may serve as functional analogs. Supporting this possibility, *in silico* protein–protein interaction predictions using FlyPredictome (Kim et al., 2024), based on AlphaFold-Multimer modeling and interface confidence scoring, indicate a potential interaction between RH7 and Gαi.

If RH7 indeed couples to a G_α_i family member, this would represent a notable exception among *Drosophila* rhodopsins by engaging an inhibitory G protein pathway. This possible reversal of signaling direction recalls the mammalian system, where melanopsin in ipRGCs activates an excitatory G_α_q pathway, while classical retinal rhodopsin signals through an inhibitory transducin (G_α_t) pathway belonging to the G_α_i family (Contreras et al., 2021; Gao et al., 2019; Koyanagi and Terakita, 2008). The use of opposing G protein pathways in these systems, despite their evolutionary distance, may reflect a conserved principle that links photoreceptor polarity to downstream signaling.

### 5.3 RH7 in the optic lobe: A modulator of early visual processing

*Rh7* is broadly expressed in the optic lobe, particularly in the lamina and medulla, which are key regions for early visual processing. These areas receive input from outer photoreceptors and are involved in contrast and motion detection. We found strong *Rh7* expression in L5 lamina neurons and Mi1 neurons of the medulla. L5 neurons pass signals from outer photoreceptors to the medulla and show strong contrast adaptation (Matulis et al., 2020). Mi1 neurons contribute to motion and contrast processing (Behnia et al., 2014; Matulis et al., 2020). The expression of *Rh7* in Mi1 neurons may thus suggest a potential role in modulating contrast-related signals, although the functional connection remains to be directly tested. These computations rely on changes in light intensity over space and time, linking the functional properties of L5 and Mi1 neurons to general luminance-based visual processing.

Our behavioral data support a role for RH7 in modulating these visual functions. Flies lacking *Rh7* show reduced optomotor responses, suggesting they are less sensitive to moving visual stimuli. In contrast, the Darkfly strain, which expresses a truncated variant of RH7, shows an increased response in the same assay (Figure 5). Rescue experiments confirm that this effect depends on *Rh7*: expression of full-length RH7 in *Rh7^0^* mutants restores normal behavior, while the truncated form of the Darkfly enhances the optomotor response. These results indicate that RH7 influences visual processing in the optic lobe. The enhanced behavior seen with the Darkfly variant suggests that changes in RH7 structure can affect its function. This possibility is discussed further in section 5.6.

### 5.4 RH7 in circadian regulation and behavioral flexibility

Fly activity is shaped by both the circadian clock and external light cues, which are detected by specialized photoreceptors (Helfrich-Förster, 2017; Rieger et al., 2003). In natural environments, these cues can vary considerably. The observation that many RH7-containing species inhabit sun-exposed environments suggests that RH7 may contribute to light sensitivity and appropriate behavioral responses under such conditions. This possibility is discussed further in the next section. In *Drosophila*, this is reflected in circadian activity patterns, to which RH7 contributes. Under standard LD 12:12 conditions, *Drosophila melanogaster* displays two peaks of locomotor activity in the morning and evening separated by a midday siesta (Helfrich-Förster, 2000). This rhythm is controlled by the circadian clock and its associated neuronal network. Flies experience two rest phases: a pronounced one at night and a shorter one during the siesta, both primarily regulated by dorsal clock neurons (DNs) (Guo et al., 2018). The DN_1_ subclass plays a key sleep-promoting role and includes CRY-positive and CRY-negative neurons that may contribute differently to the two rest phases (Guo et al., 2016; Lamaze and Stanewsky, 2020).

RH7 is expressed in dorsal clock neurons, especially in CRY-negative DN_1_, with the exception of the two DN_1a_ neurons (4F). While we could confirm RH7 expression in the DNs, we did not detect it in lateral clock neurons (Figure 4B, D), as previously suggested by Ni et al. (2017). Ni et al. (2017) reported that RH7 plays a central role in circadian light adaptation and observed its expression in lateral neurons (LN_v_s). However, Senthilan et al. (2019) later found that the original antibody staining in LN_v_s was likely artifactual, and Lazopulo et al. (2019) further demonstrated that RH7’s functional effects on light-dependent behavior occur outside the LN_v_s. In addition to its role in DNs, RH7 may also influence circadian regulation via the optic lobe. Its expression in the lamina and medulla, along with possible synaptic connections between Mi1 neurons and PDF-expressing neurons (Reinhard et al., 2024), and the presence of PDF receptors in lamina neurons (Bastian et al., 2025; Davis et al., 2020), suggest a functional link between early visual processing centers and the circadian network. Rather than acting as a dominant regulator, RH7 appears to exert subtle, context-dependent effects shaped by environmental conditions and genetic background. As a result, its contribution may be masked by other photoreceptive pathways or background variation, leading to differing phenotypes across studies. Furthermore, the extent to which certain behaviors are restored may depend on how closely the *Rh7-Gal4* driver recapitulates the endogenous *Rh7* expression pattern, as some *Rh7*-expressing cells, especially in the visual system, were detected only by *in situ* hybridization (Figure 4G,4H). Together, our findings contribute to a better understanding of RH7 function in light-dependent behavior and provide a basis for future studies to further define its potential role in circadian entrainment.

### 5.5 Evolutionary retention of RH7 in sun-exposed species

The repertoire of opsins and cryptochromes varies considerably among species, reflecting lineage-specific adaptations to different light environments. Yet, a consistent evolutionary link exists between RH7 and DCRY within the light-sensing network. While DCRY is evolutionarily older and widespread across Bilateria, RH7 arose later within Panarthropoda. Consequently, many non-Panarthropoda species possess DCRY, but not RH7. In contrast, nearly all arthropod species that retain DCRY also possess RH7 (approximately 98%), indicating a strong evolutionary link. However, RH7 can exist independently of DCRY. Lineages that have lost DCRY often retain RH7, particularly in species exposed to high sunlight, especially during early developmental stages (e.g., certain sawflies and some beetles).

This pattern suggests that RH7 contributes to light sensitivity and the regulation of behavior under natural, sunlit conditions Meyerhof et al. (2024) reported altered light-dark activity in *Rh7* mutants, highlighting a modulatory role for RH7 in behavioral responses to ambient light. In our LD and wavelength assays, WT flies showed higher activity during dawn and dusk and a pronounced siesta, whereas *Rh7^0^* mutants displayed increased daytime activity and a reduced siesta. Avoiding activities at midday, when light intensity is at its highest under natural conditions, is consistent with a protective role of RH7, as suggested by Lazopulo et al. (2019) under high-intensity blue light. The moderate light intensities in our experiments were likely below the threshold for strong avoidance. Yet they still revealed behavioral differences between WT and *Rh7^0^* flies. These results support a role for RH7 in mediating protective behavioral responses to environmental light. While its precise functional contribution outside of *Drosophila* remains to be determined, the widespread retention of RH7 across diverse light-exposed species highlights its likely adaptive and conserved role in detecting and responding to environmental light.

### 5.6 RH7 is essential for locomotor activity during darkness

Although activity during darkness is regulated by the circadian clock, the startle response to light-off is independent of it. Dark-pulse experiments show that both WT controls and *per^0^* mutants, which lack a functional circadian clock, exhibit a strong startle response, consistent with the absence of diel modulation of the response in *per^0^*mutants (Figure 3B). While the circadian clock seems to modulate the magnitude of the response, it is not essential for its occurrence (Figure 3B). Moreover, this phenotype persists in flies rendered arrhythmic during the 4-hour assay. The critical role of RH7 is supported by trans-heterozygous combinations with *Rh7^MiMIC^*, which show the same phenotype as *Rh7^0^*mutants, and by rescue experiments restoring normal behavior (Figure 3–figure supplement 2B).

The Darkfly strain, bred in complete darkness for many generations, provides an intriguing case for RH7 function in sustained darkness. It carries an RH7 variant with a 21-amino acid deletion in the C-terminal tail, a region rich in predicted phosphorylation sites. Structural modeling based on AlphaFold predicts an interaction between this tail and the receptor core, specifically the unique ICL3 loop of RH7 (Figure 1A), which lacks the QAKKMNV motif found in other *Drosophila* rhodopsins. Such intramolecular interactions are known to mediate autoinhibition in GPCRs, where the C-terminal tail stabilizes an inactive conformation that shifts to an active state upon receptor stimulation (Maggio et al., 2023).

We hypothesize that the deletion in the Darkfly variant weakens this autoinhibitory interaction and thus facilitates constitutive activation (Figure 6B). This is supported by the observation that deleting the final 21 amino acids increases startle response and optomotor response, approaching levels seen in Darkfly flies (Figure 5). However, the unnormalized locomotor activity data (Figure 5 –figure supplement 1A) reveal generally elevated locomotor activity in the Darkfly strain, likely reflecting additional genetic changes accumulated during long-term maintenance in constant darkness (Izutsu et al., 2012), which may also influence neural development or synaptic connectivity. For example, nonsense mutations in *HisCl1* and *Vha100-1* could alter visual processing and neural signaling (Alejevski et al., 2019; Williamson et al., 2010), while missense mutations in *DopR*, *bgm*, and development-associated transcription factors *Sox21b* could modulate activity, sleep, and neural development (Jiang et al., 2016; McKimmie et al., 2005; Min and Benzer, 1999). Taken together, these findings highlight the pivotal role of RH7 in modulating behavior in darkness and demonstrate that structural changes can fine-tune RH7 signaling to adapt to environmental conditions.

### 5.7 Parallels and divergence between *Drosophila* RH7 and other non-canonical Opsins

This study highlights significant similarities between RH7 and other noncanonical rhodopsins, particularly mammalian melanopsin (OPN4). Both RH7 and melanopsin are expressed in neurons downstream of classical photoreceptors, with RH7 in lamina and medulla neurons of the optic lobe and melanopsin in the ipRGCs, suggesting analogous roles in processing visual and circadian information. Functionally, both opsins contribute to circadian regulation as well as contrast and motion detection. Structurally, they share elongated N- and C-terminal regions that are intermediate in length relative to canonical rhodopsins and typical GPCRs. These extended regions harbor multiple phosphorylation sites that are likely crucial for their unique signaling mechanisms. In addition, we identified five conserved sequence motifs shared between RH7 and melanopsins, including two in the C-terminal region and others within the transmembrane core, one of which extends into the conserved DRY motif in the second intracellular loop (ICL2) (see Figure 6 –figure supplement 1A, 1B).

Furthermore, RH7 and melanopsin activate phototransduction cascades distinct from those of classical visual opsins, potentially producing opposing effects. This reflects an evolutionary adaptation that supports diverse functions in both visual and nonvisual photoreception. However, while melanopsin expression in mammals is largely confined to the visual system, *Drosophila* RH7 appears to have acquired additional functions in the brain. This divergence may reflect evolutionary adaptations to differing anatomical constraints: the small, light-permeable brain of *Drosophila* may permit direct light sensing beyond the eyes, whereas in mammals, the evolution of a larger brain enclosed by a light-impermeable skull may have limited such extraocular photoreception to specialized retinal pathways.

Although melanopsin is predominantly expressed in retinal ganglion cells, it has also been detected in various brain regions (Aviles-Trigueros et al., 2012; Nissilä et al., 2017). Additionally, other non-canonical opsins such as encephalopsin and neuropsin are present in the human brain, with their functional roles still under investigation (Buhr et al., 2015; Nissilä et al., 2012; Upton et al., 2022, 2021). In this context, the expression of RH7 in the *Drosophila* brain, outside of clock neurons, may reflect similar non-visual roles. While its brain expression is relatively weak compared to its prominent presence in the optic lobes, it is consistently observed using both *Rh7*-*Gal4* reporters and *in situ* hybridization. The identity of the non-clock neuronal populations expressing *Rh7* remains to be determined, and further work, such as double labeling with cell-type-specific markers, will be necessary to clarify this. In these neurons, RH7 might contribute to higher-order visual processing, sensorimotor control, general arousal, or aspects of neuronal development.

These findings open new possibilities to study how noncanonical opsins like RH7 contribute to complex brain functions beyond just sensing light. Understanding the roles of RH7 can give important clues about how light-sensing systems evolved and how they work together with brain circuits to control behavior. Future research using genetics, anatomy, and physiology will be important to fully understand all the functions of RH7 in *Drosophila* and other animals.

## 6 Conclusion

Our study provides a key insight into the unresolved functional diversity of opsins. Unlike canonical rhodopsins involved in image formation, RH7 acts as a fundamental photoreceptor mediating simple light-off signals to diverse neuronal populations. As an evolutionarily ancient opsin, RH7 likely represents a primordial photoreceptive system predating the specialization of complex visual pathways. Structurally and functionally, RH7 exhibits characteristic features of G protein-coupled receptors (GPCRs), positioning it as an evolutionary intermediate between ancestral GPCRs and specialized visual opsins. This is reflected in its atypical sequence motifs, extended terminal domains, and distinct signaling properties, which collectively blur the distinction between classical visual and non-visual photoreception.

Functionally, RH7 modulates early visual processing in the lamina and might contribute to circadian regulation in dorsal clock neurons. Its phylogenetic and mechanistic parallels to mammalian melanopsin further support its role as a GPCR-like photoreceptor integrating environmental light cues with intracellular signaling cascades to regulate physiology and behavior. Notably, many RH7-retaining species, with or without DCRY, inhabit environments with high light exposure. This ecological context may indicate a potential adaptive role for RH7 in mediating light sensitivity in naturally sun-exposed environments. In summary, RH7 exemplifies how ancient opsins have diversified to couple photoreception with GPCR-mediated signaling beyond image formation. Its expression in brain regions highlights a conserved evolutionary strategy: the integration of light sensitivity into central neural circuits, underscoring the deep-rooted link between sensory and neurodevelopmental architectures.

## 7 Resource availability

All data supporting this study are available within the main text or the Supplemental Information. Additional details can be obtained from the corresponding author upon request.

## 8 Acknowledgements

We thank Kathrin Sauter, Christina Brunner, Amelie Brönner, and Marina Müller for technical assistance. We also acknowledge Maria Gallant and Barbara Mühlbauer for their support, and Giulia Manoli for help with data analysis. We are grateful to Naoyuki Fuse for providing Darkfly, to E. Kostenis and A. Inoue for the HEK293ΔG7 cell line, and to Martin Göpfert and Marion Silies for valuable discussions. This work was supported by the German Research Foundation (DFG), SE 2320/1-1, SO 1337/2-2, and the DFG Heisenberg program SO 1337/6-1.

## 9 Author contributions

V.K. performed fly activity assays, optomotor experiments, generated UAS-construct lines, and carried out MCFO staining and qPCR analyses. N.R. performed immunohistochemistry and in situ hybridizations. H.H. generated the *Rh7-Gal4* line. A.M. contributed additional fly activity assays. D.R. established experimental setups and contributed to experimental design. P.S. carried out the GsX assay. C.H.-F. contributed to conceptual discussions, provided critical input throughout the project, and performed dissections for qPCR. P.R.S. designed the experiments, coordinated the study, analyzed experimental results, performed phylogenetic analyses, and wrote the manuscript. All authors reviewed and approved the final manuscript.

## 11 Declaration of generative AI and AI-assisted technologies in the writing process

During the preparation of this work, the authors used DeepL and ChatGPT in order to improve the readability and language of the manuscript. After using these tools, the authors reviewed and edited the content as needed and take full responsibility for the content of the published article.

