## Supplementary material for "RHODOPSIN 7: An ancestral non-canonical photoreceptor shaping light-responsive behavior": Figure 3 - supplementary figure 1

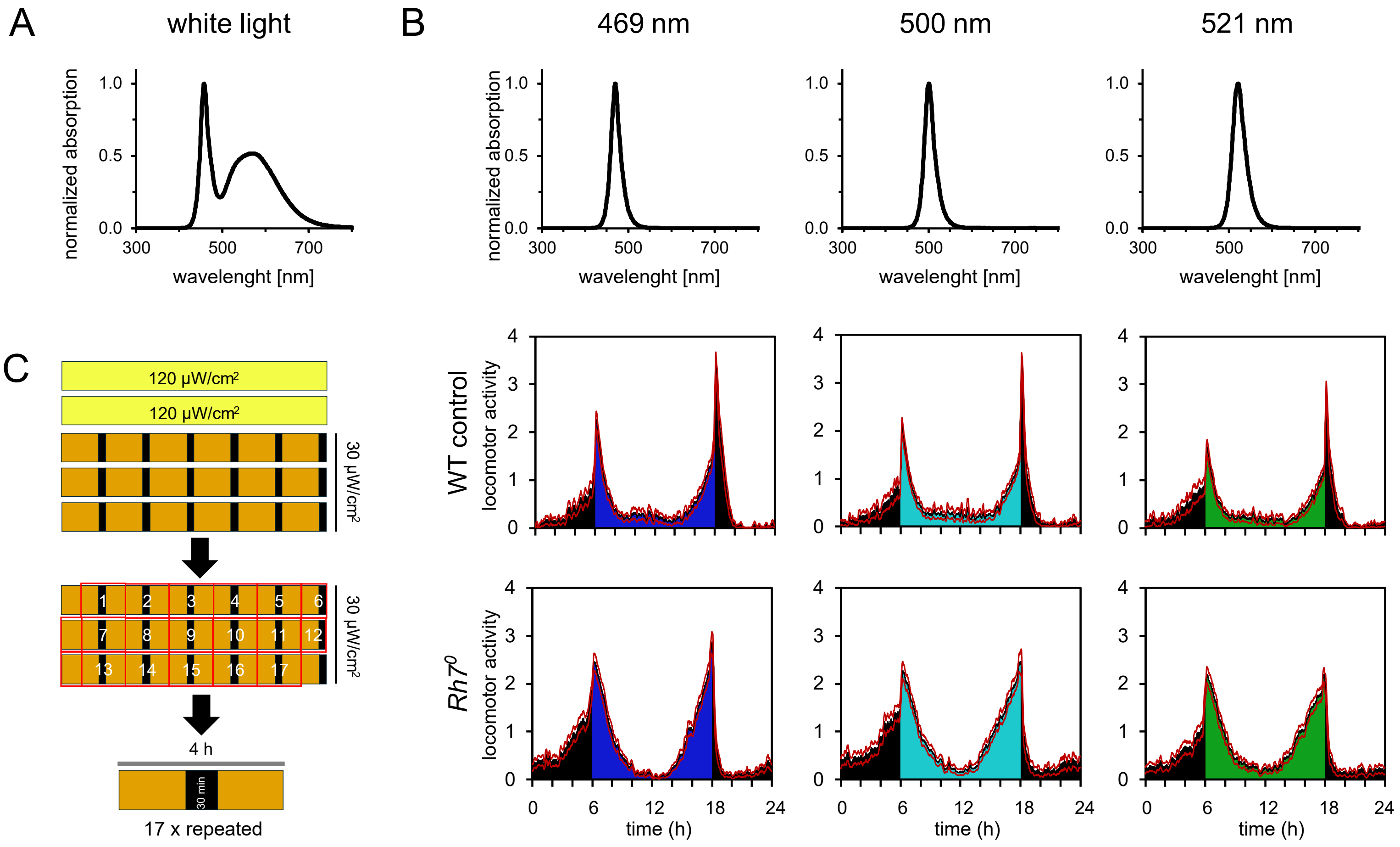


**Figure 3 – figure supplement 1 Spectral characteristics of the light stimuli and experimental design**.

**(A)** Absorption spectrum of the white light used in the experiments. **(B)** Absorption spectra of the 469 nm, 500 nm, and 521 nm light stimuli (top), together with the average locomotor activity profiles of WT control flies (middle) and *Rh7*$^0$ mutant flies (bottom) under standard conditions (12 h light (30 µW/cm²):12 h dark, 25 °C). **(C)** Schematic representation of the 4 h assay protocol.
