## Supplementary material for "RHODOPSIN 7: An ancestral non-canonical photoreceptor shaping light-responsive behavior": Figure 3 - supplementary figure 2

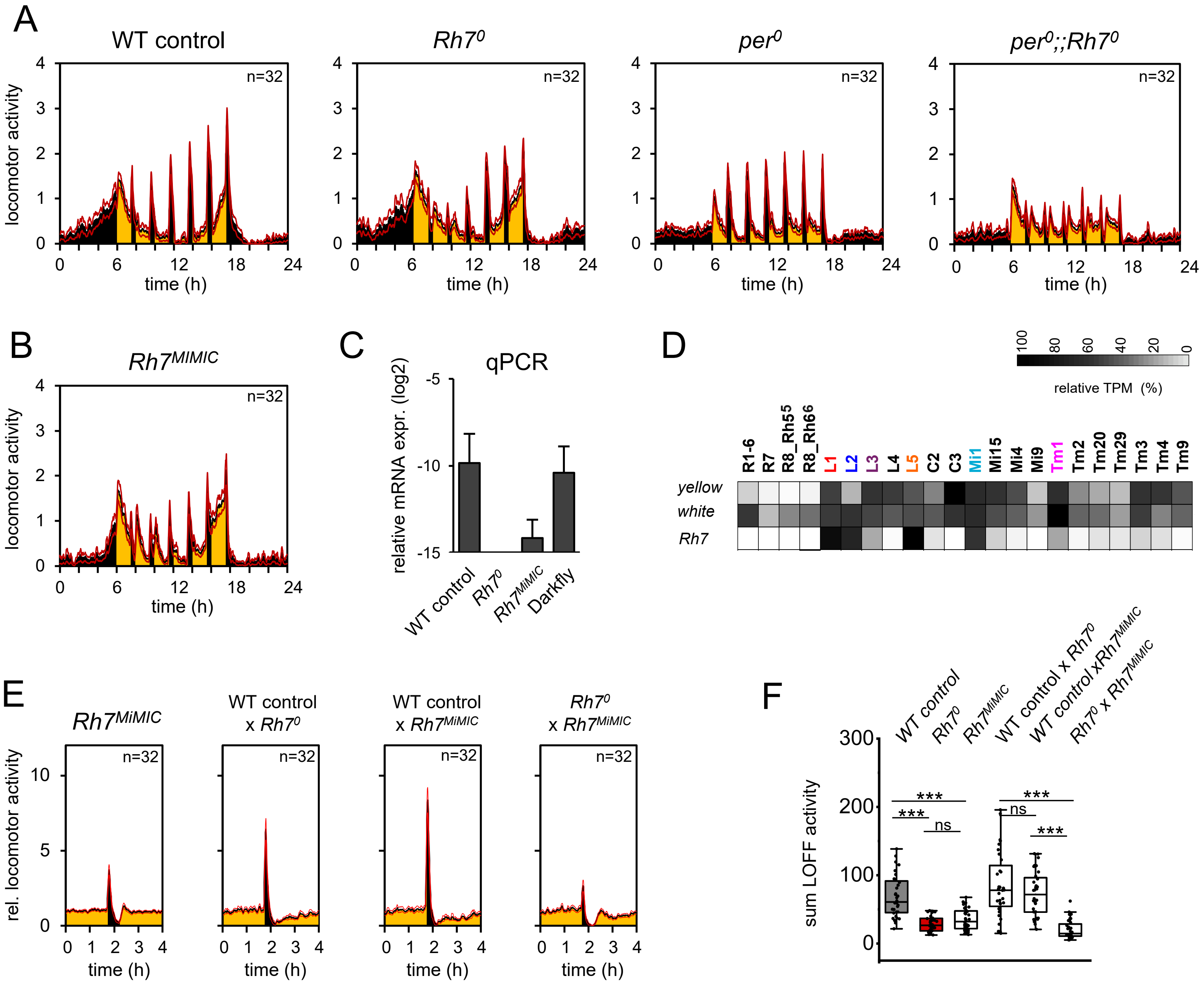


**Figure 3– figure supplement 2 Behavioral and gene expression analysis of different mutant and control lines.**

**(A)** Non-normalized average locomotor activity profiles of WT control, *$Rh7^0$*, *$per^0$*, and *$per^0$;;$Rh7^0$* flies under standard conditions (12 h light (30 µW/cm²):12 h dark, 25 °C) with dark pulses. **(B)** Non-normalized average locomotor activity profile of the *Rh7MiMIC* line under LD 12:12 conditions with dark pulses. The activity profile is similar to that of the *$Rh7^0$* mutant. **(C)** qPCR analysis of *Rh7* mRNA expression in whole heads of WT control, *$Rh7^0$*, *$Rh7^{MiMIC}$*, and Darkfly flies. *Rh7* transcript levels were comparable between WT control and Darkfly flies, absent in the *$Rh7^0$* mutant, and reduced in the *$Rh7^{MiMIC}$* line. **(D)** Heatmap of relative gene expression across *Drosophila* optic lobe cell types. Expression of *Rh7*, *yellow*, and *white* is shown. TPM values were obtained by TAPIN sequencing (Davis et al., 2020) and normalized independently for each gene (100% = maximum TPM per gene). The scale ranges from white (0%) to black (100%). **(E)** Startle responses in the 4-hour assay for *$Rh7^{MiMIC}$*, *$Rh7^0$/$Rh7^{MiMIC}$* trans-heterozygotes, and the corresponding crosses to WT controls. *$Rh7^{MiMIC}$* flies and *$Rh7^0$/$Rh7^{MiMIC}$* trans-heterozygotes exhibited strongly reduced startle responses, whereas the corresponding WT control crosses showed normal responses. **(F)** Box plots showing the summed dark activity of the fly lines analyzed in (E), (***$p < 0.001$, Mann-Whitney U-Test, BH correction, R-studio).
