## Supplementary material for "RHODOPSIN 7: An ancestral non-canonical photoreceptor shaping light-responsive behavior": Figure 4 - supplementary figure 1

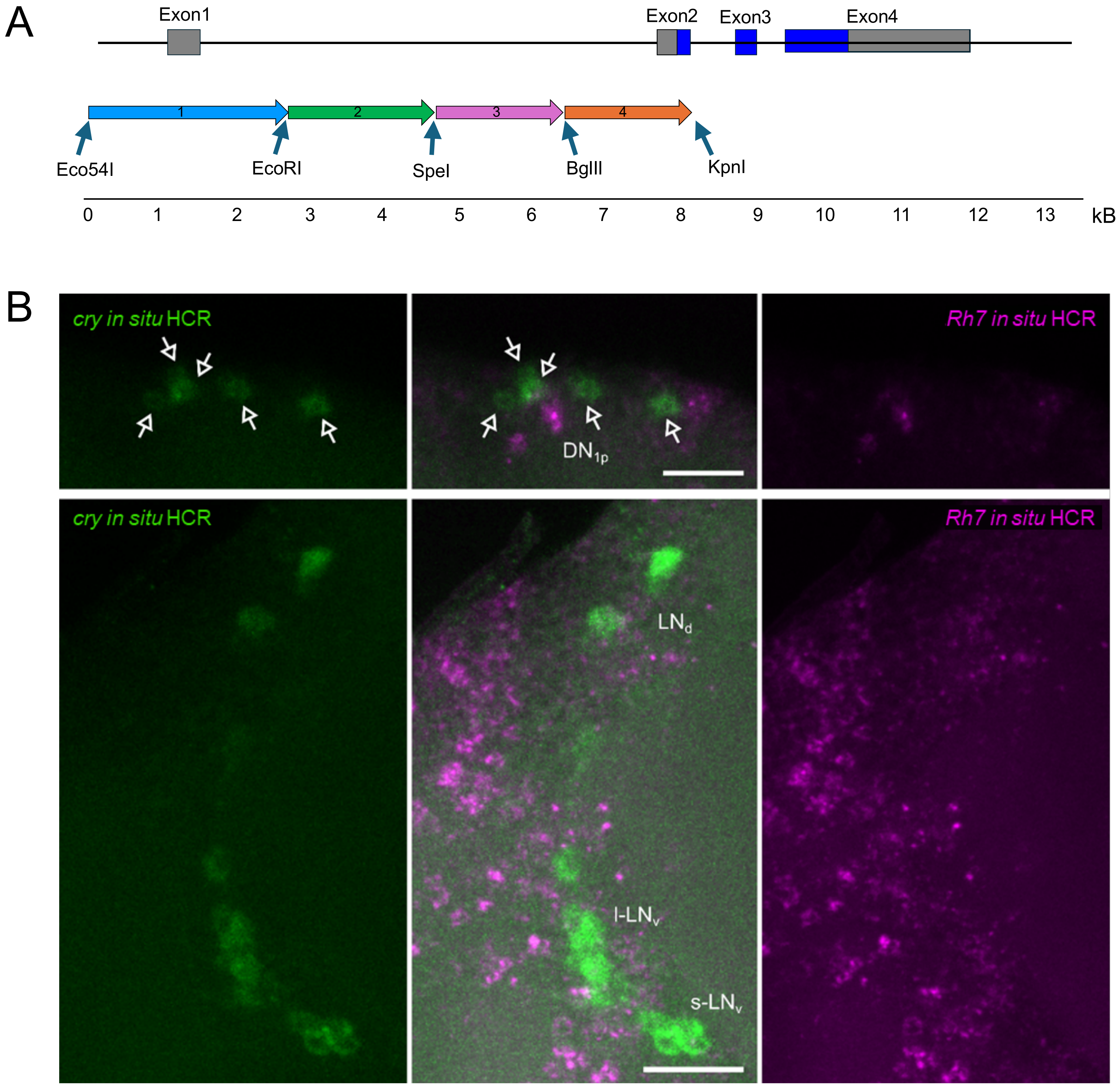


**Figure 4– figure supplement 1 Generation of the *Rh7-Gal4* driver and *Rh7* and *cry* expression analysis.**

**(A)** Schematic overview of the cloning strategy used to generate the *Rh7-Gal4* driver line. An ~8 kb genomic fragment containing the *Rh7* promoter region, including exons 1 and 2, was assembled in four sequential cloning steps.
**(B)** Double *in situ* hybridization for *Rh7* and *dcry* (*cry*) in brains of WT control flies.
