## Supplementary material for "RHODOPSIN 7: An ancestral non-canonical photoreceptor shaping light-responsive behavior": Figure 5 - supplementary figure 1

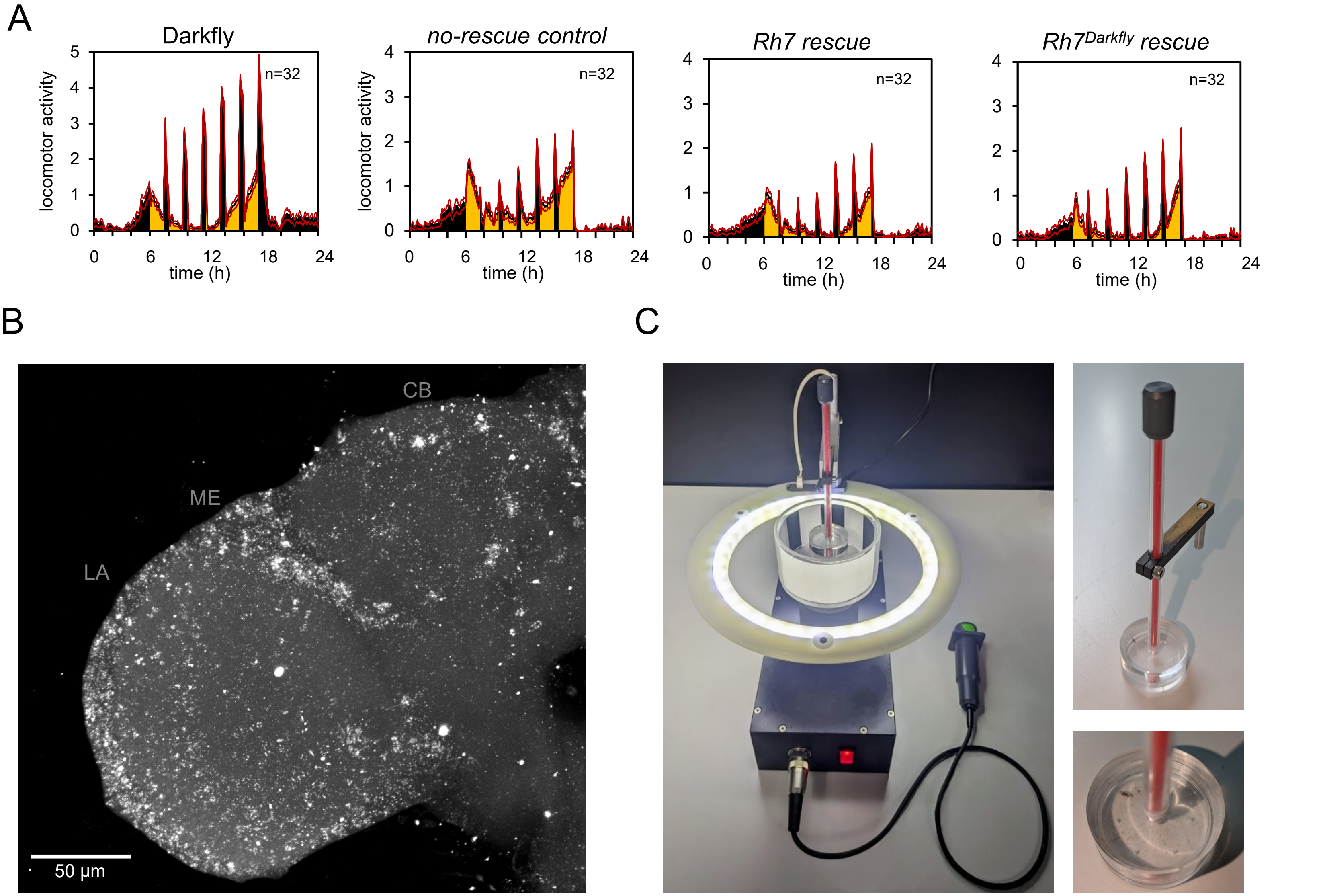


**Figure 5 – figure supplement 1. *Rh7* rescue lines, *Rh7* expression in Darkfly, and the optomotor assay.**

**(A)** Non-normalized average locomotor activity profiles of Darkfly, the no-rescue control (*Rh7-Gal4; Rh7$^{0}$ > UAS-empty; Rh7$^{0}$*), the *Rh7* rescue (*Rh7-Gal4; Rh7$^{0}$ > UAS-Rh7; Rh7$^{0}$*), and the *Rh7$^{Darkfly}$* rescue (*Rh7-Gal4; Rh7$^{0}$ > UAS-Rh7$^{Darkfly}$; Rh7$^{0}$*) under standard conditions (12 h light (30 µW/cm²):12 h dark, 25 °C) with dark pulses. **(B)** RNA *in situ* hybridization for *Rh7* in Darkfly brains, showing similar expression patterns like WT controls. **(C)** Experimental setup of the optomotor assay. A video of the assay is provided as Supplementary Video 1.
