## Supplementary material for "RHODOPSIN 7: An ancestral non-canonical photoreceptor shaping light-responsive behavior": Figure 6 - supplementary figure 1

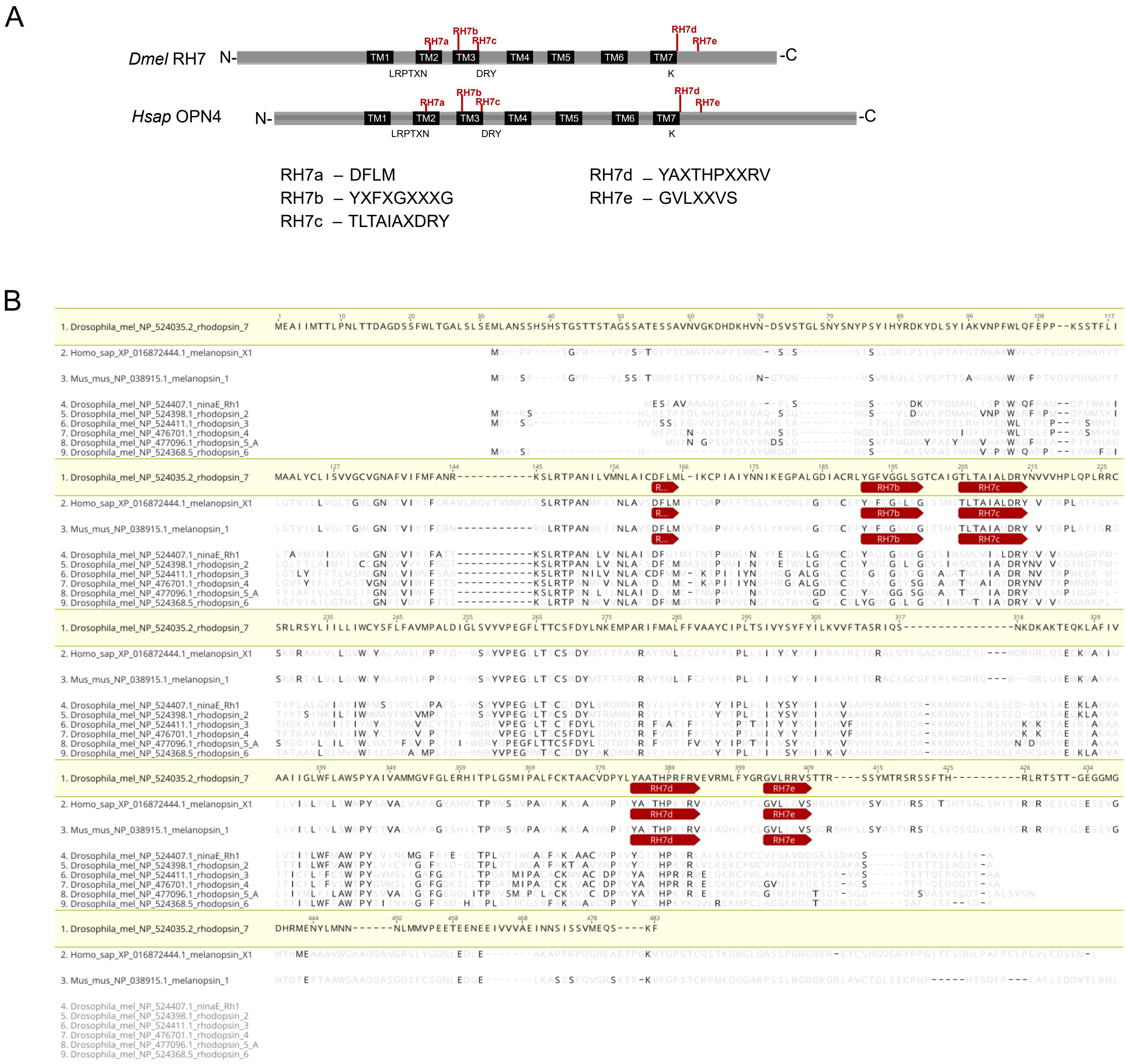


**Figure 6– figure supplement 1 Sequence comparison of RH7/MELANOPSIN protein sequences.**

**(A)** Alignment of the protein sequences of *Drosophila melanogaster* RHODOPSIN 7 (*Dmel* RH7) and *Homo sapiens* MELANOPSIN (*Hsap* OPN4), identifying five conserved motifs shared between the two proteins. **(B)** Protein sequence alignment of *Drosophila melanogaster* rhodopsins and melanopsin (OPN4) sequences from *Homo sapiens* and *Mus musculus.* Red arrows indicate the five newly identified conserved motifs.
